# Phosphorus-laden Mg/Fe Layered Double Hydroxide Dispersed on Douglas fir Biochar as a Controlled Release Fertilizer and its effect on the growth of bush beans (Phaseolus vulgaris)

**DOI:** 10.64898/2026.05.22.727001

**Authors:** Tajinder Singh, Prashan M. Rodrigo, Ryan A. Folk, Jagmandeep Dhillon, Jac J. Varco, Todd E. Mlsna

## Abstract

Many agricultural soils are deficient in key macronutrients needed for healthy plant development. Relying on highly water-soluble commercial fertilizers for long durations can be costly and environmentally harmful. This study investigates a phosphorus-loaded Mg/Fe layered double hydroxide (LDH) dispersed on Douglas fir biochar (Mg/Fe-LDH biochar) as a controlled-release fertilizer and evaluates its impact on bush bean (*Phaseolus vulgaris L*.) growth. Emphasizing sustainability, the work integrates controlled-release fertilizers, biochar, and LDH modification to enhance nutrient use efficiency and mitigate environmental runoff. Mg/Fe-LDH was directly synthesized on biochar via a co-precipitation approach, loaded the composite with phosphate by anion exchange, and characterized the material using elemental analysis, N_2_ Brunauer-Emmett-Teller (BET) determinations surface area analysis, and x-ray photoelectron spectroscopy to confirm successful LDH modification on Douglas fir biochar, and high surface area with accessible active sites. The synthesis yielded a stable P-Mg/Fe-LDH biochar with enhanced dispersibility and phosphate-buffering capacity, enabling controlled-release fertilization. In greenhouse experiments, bush beans grown with the P-Mg/Fe-LDH biochar exhibited improved growth metrics, including increased yield (beans fresh weight of 31.7 g), biomass (plant dry weight of 6.3 g), plant height (32.8 cm), and improved nutrient uptakes (1.88 mg (P) g^-1^) at 100.88 kg (P_2_O_5_) ha^-1^ compared with unfertilized controls and conventional P fertilizers, indicating efficient, controlled-release phosphate delivery and sustained nutrient availability. The results demonstrate that integrating LDH-modified biochar can enhance P uptake and plant growth while reducing leaching losses. Overall, this study highlights the strategic significance of combining biochar, layered double hydroxides, and controlled-release formulations to advance sustainable nutrient management and improve crop performance in agroecosystems. The findings offer a promising pathway for environmentally conscious fertilizer design and soil amendment strategies that align with global goals for resource efficiency and food security.

**Graphical Abstract:** 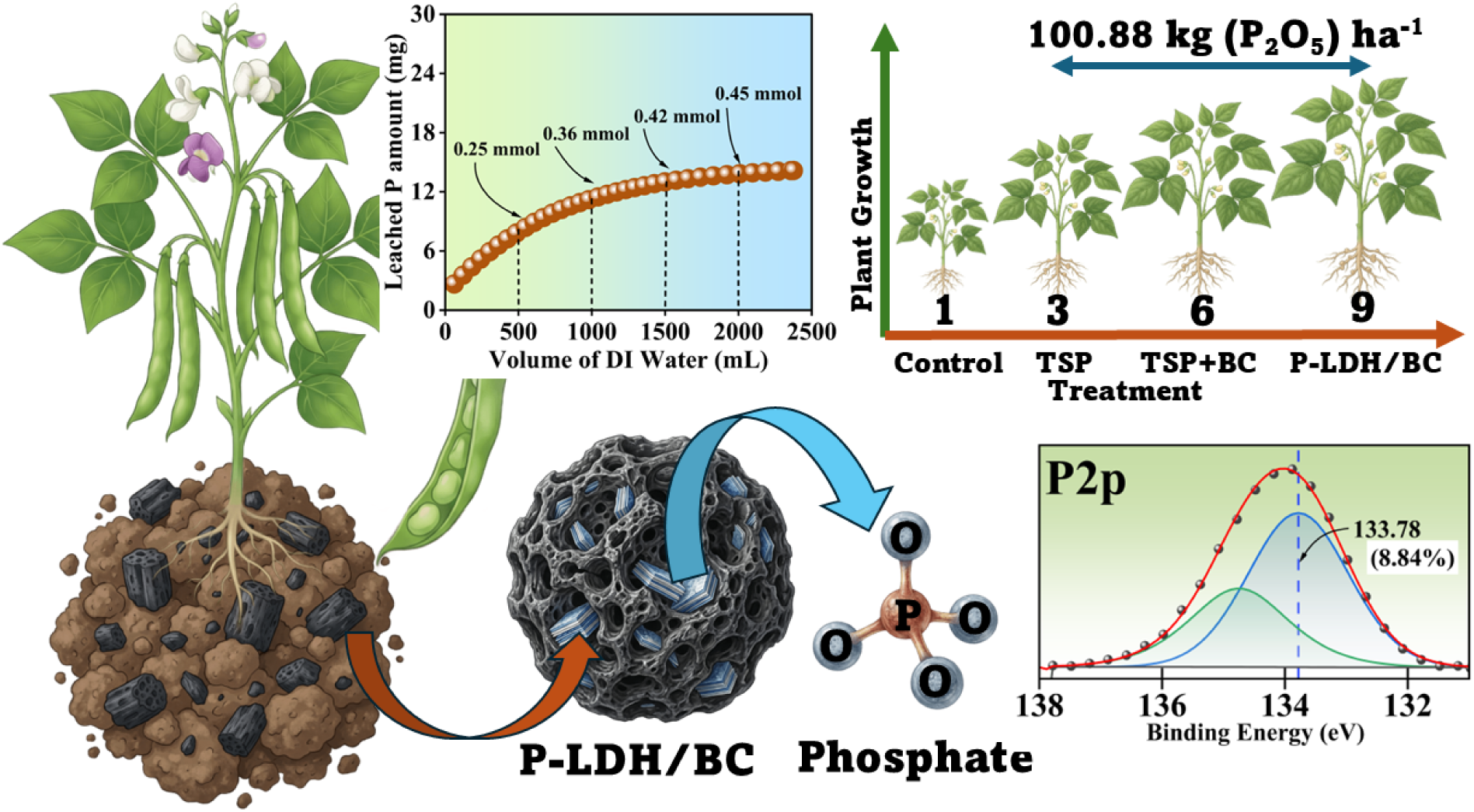

## 1. Introduction

Agriculture has long been recognized as a significant contributor to water pollution via nonpoint sources (Guo et al., 2014). Recent U.S. Environmental Protection Agency national water-quality inventories noted that nonpoint sources ranked in five of the six top spots in their contribution to water-quality-related impairments of rivers and streams in the U.S. Moreover, agriculture-related activities were consistently identified as the top point sources of water-quality impairments across the U.S. Phosphorus (P) and nitrogen (N) were reported as the most widespread stressors in lakes, ponds, and reservoirs, with 21% of the nation’s lakes being hypereutrophic.

Phosphorus is known as the main stressor in coastal waters in the U.S. Nutrient runoff is a major nonpoint source of water pollution due to either mis-application (e.g., timing and rate) and weather events causing leaching or runoff of soluble nutrients derived fertilizers and manures (Paulson et al., 2010). Recent advancements in soil testing and informed fertilizer amendments help address these issues (Micha et al., 2023), but a major global shift to responsible nutrient management is yet to be realized. Reactions of nutrient elements in soil depend on the union of biotic transformations and abiotic chemical interactions. Adsorption reactions depend heavily on ionic charge and soil adsorption capacity, thus individual management strategies may be necessary (Hodges, 2010), but difficult to implement. Phosphorus is unique in its ability to bind to soil particles, forming complexes that vary from labile to non-labile. Adsorption and or chemically precipitated complexes may slowly dissolve into their bioavailable forms depending on soil characteristics: pH, organic content, moisture, etc (Schröder et al., 2011). To avoid relatively low bioavailable P levels early in the growing season due to cold temperatures slowing diffusion processes, growers apply highly soluble inorganic P fertilizers including polyphosphate solutions during the early growing season, especially for responsive crops such as corn (*Zea Mays*) and wheat (*Triticum Aestivum*). The efficiency of P applied to agricultural fields remains low, with only 10-15% of applied P taken up by crops in field studies (Dhillon et al., 2017), while the remainder is adsorbed or precipitated in the soil with a resulting lower availability. Fertilized agricultural lands contain significant reserves of adsorbed phosphorus, enough to support future crops for decades (Roberts et al., 2015). Chemically adsorbed forms of soil P can enter waterways through soil erosion, with subsequent desorption of adsorbed soil P due to equilibrium shifts under more dilute conditions. The background nutrient concentrations in streams draining predominantly agricultural watersheds are nine times higher than in those draining from predominantly forested watersheds (Brown et al., 2012). As surface waters are often P limited, the increased nutrient loading from streams can result in eutrophication and cause harmful algal blooms in aquatic systems (Brown et al., 2012). Inefficient P use and current consumption patterns along with costly purchased of chemical fertilizers put undue economic pressures on growers and sustainable crop production (Childers et al., 2011; Daneshgar, Buttafava, et al., 2018). There is a need for transformational changes in resource efficiency, as P has been classified as an essential critical raw material (Golroudbary et al., 2019; Scholz et al., 2013). Moreover, even though there is no imminent predicted shortage of phosphate rock, with global reserves amounting to over 300 billion metric tons, the current technology and economics deem a significant portion unavailable for extraction in the near future (Daneshgar, Callegari, et al., 2018). Concentrated sources of extractable phosphate rocks are limited and distributed disproportionately across a handful of countries. The global production from high-grade phosphate rocks, the current primary source of P, is estimated to surpass demand between 2035 and 2090 (Jacobs et al., 2017). The looming production peak threatens global food security, and the challenges related to supplies will be unprecedented as P is an indispensable mineral for agriculture (Jacobs et al., 2017). Low phosphate use efficiency, environmental concerns, and potential production peaks thus threaten global food security and motivate sustainable use and investment in P recovery and recycling (Scholz et al., 2019).

Layered double hydroxides (LDH) are candidates for recovering and recycling phosphates from wastewater sources (Keyikoglu et al., 2022). The phosphate adsorbed on LDH has a secondary fertilizer benefit in that it acts as a buffer for soluble P in agricultural systems, thus preventing its conversion to non-labile forms in soil and ensuring longer availablity for plant uptake (Bernardo et al., 2018). LDHs have a general chemical formula of 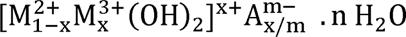 where M^2+^ and M^3+^ represent a di- and trivalent cation, respectively, A^m-^ is an anion interspersed with a load m, x is the di- and trivalent cation ratio, n is the moles of water (Wu et al., 2018). They can be synthesized in the laboratory with a standard co-precipitation method where aqueous divalent and trivalent metallic salt solutions are mixed with the desired anion, followed by a pH adjustment (Edañol et al., 2020). The interlayer anions are the guest layer, which other anions can exchange in the water, while weak hydrogen bonds from the surrounding water molecules hold the structure together (Keyikoglu et al., 2022). The metal hydroxides and their environmental characteristics enable the adsorbent properties of the LDH to exchange phosphates and other contaminants for the interspersed anions (Keyikoglu et al., 2022; Tang et al., 2022). The adsorbed phosphates can be recovered and used as controlled-release fertilizers (Bernardo et al., 2018). Studies on structural properties, adsorption, and desorption mechanisms of LDH support their application in phosphate recovery (Edañol et al., 2020; Keyikoglu et al., 2022; Tang et al., 2022). The application of phosphorus-loaded LDH in greenhouse and field experiments requires further investigation to establish its potential as a controlled-release fertilizer, which we aim to achieve with our study. Applications of LDH in sorption systems face critical structural challenges like particle clustering, low mechanical strength, and aggregation (Henrist et al., 2003). These challenges reduce LDH adsorption efficiency and limit their application for P recycling. Exfoliating the LDH into a few-layered nanosheet helps reduce aggregation and increase surface area, exposed sites, and dispersibility (Peng et al., 2021). More recently, in-situ dispersion of small LDH particles on a larger, high surface area biochar has also been used to prevent particle aggregation, increase sorption capacity and rates, and provide extra surface area to remove additional sorbates (Rahman et al., 2021).

Biochars are usually by-products of wood pyrolysis and are relatively inexpensive, making them desirable as dispersion media for precipitating LDH nanoparticles. Biochars are sorbents for metal cations and organic sorbents (Mohan et al., 2014; Rodrigo & Kommalapati, 2025). Biochar amendments to soil also increase organic content, enhance water and nutrient retention, and help maintain a soil profile that supports agriculture (Rodrigo, Varco, et al., 2025). The Douglas fir biochar used in this study is a cheap, commercially available, carbon-rich adsorbent matrix with high surface area and pore volume available for LDH dispersion (Arwenyo et al., 2024; Rahman et al., 2021). It is a by-product of wet wood fast pyrolysis under low oxygen conditions in an updraft gasifier at 900-1000 °C (1 -10 s). Aqueous Magnesium and Iron salts were selected as the di and trivalent metal cations, respectively, mixed in a 2:1 molar ratio to obtain a high surface area Mg/Fe-LDH complex dispersed on the Douglas fir biochar (Mg/Fe-LDH biochar). The LDH-biochar complex was loaded with phosphates via anion exchange in an aqueous medium to obtain P-Mg/Fe-LDH biochar. The P-Mg/Fe-LDH biochar was tested as a controlled-release fertilizer in a controlled greenhouse experiment on common bush type beans (*Phaseolus vulgaris L.*). Bush type beans are nodulating legumes and host nitrogen-fixing bacteria which satifies a portion of the nitrogen requirement, resulting in P being a main focus as the most limiting nutrient for plant growth. Common garden bush beans are are easy to grow, have a relatively short days to maturity from emergence (45-60 days), and are limited in stature and less prone to rootbound conditions under our proposed growth cycle., The overall objective of this study was to test the effects of P amendments derived from LDH-modified biochar on the growth and nutrition of common garden bush beans.

## 2. Methodology

### 2.1 Chemicals and Materials

All chemicals (MgCl_2_⋅6H_2_O, FeCl_3_, NaH_2_PO_4_⋅H_2_O, and NaCl) used, unless otherwise noted, were analytical grade and supplied by Sigma-Aldrich. Douglas fir biochar (BC) was provided by Biochar Supreme, Everson, WA, for this study. BC is a by-product from wet wood gasification of waste Douglas fir timber chipped (∼3 in.) and auger-fed. The green wood was fed to an air-fed updraft gasifier with a 1-10 s residence time at 900 – 1000 °C. The large biochar particles (∼2 cm) were washed with 1 L deionized (DI) water, vacuum filtered, and oven-dried for 24 h at 105 °C. The elemental analysis of the filtrate is reported in Table S1. The dried biochar was ground, sieved to 250 - 420 µm particle size, and stored in closed vessels for characterization and later use.

### 2.2 Preparation of layered double hydroxide (LDH) dispersed on Biochar (Mg/Fe-LDH/BC)

LDH/BC was synthesized in a 2:1 molar ratio of Mg^2+^: Fe^3+^ in an alkaline medium and dispersed in situ on biochar. A 25.0 g of dried biochar (250-420 μm) was mixed with FeCl_3_ (8.97 g) and MgCl_2_⋅6H_2_O (10.55 g) in 300 mL of deionized water to obtain a slurry. The pH of the slurry was increased to 13 using 4 mol L^-1^ NaOH and stirred (200 rpm) for 24 h at 25 °C. The prepared LDH biochar was vacuum filtered and washed with 100 mL pure ethanol, followed by a 200 mL deionized water wash. LDH/BC was oven-dried for 24 h at 105 °C. The final weight of LDH/BC was 41.08 g and stored in a closed polypropylene container.

### 2.3 Preparation of P-Mg/Fe-LDH/BC

A 5 g L^-1^ P stock solution was prepared by dissolving 22.28 g of NaH_2_PO_4_⋅H_2_O in 1 L of deionized water and adjusting pH to 7 with 4 mol L^-1^ NaOH solution. 30.00 g of the prepared LDH/BC was mixed with 750 mL of the 5 g L^-1^ P solution and stirred (200 rpm) for 3 h in an orbital shaker to obtain phosphorus-laden LDH biochar (P-LDH/BC). The P-LDH/BC was vacuum-filtered and oven-dried for 24 h at 105 °C. The final weight was 24.64 g and stored in a glass container. The loss of 5.36 g is attributed to partial dissolution of soluble minerals from the Douglas fir biochar, detachment of loosely bound Mg/Fe-LDH phases during phosphorous loading and filtration, and minor solid loss during vacuum filtration.

### 2.4 Characterization of BC, Mg/Fe-LDH/BC, and P-Mg/Fe-LDH/BC

The BC, LDH/BC, and P-LDH/BC were characterized by point of zero charge (PZC), Electrical Conductivity (EC), X-ray photoelectron spectroscopy (XPS), surface area measurement, and elemental composition. Leachate and column studies estimated the amount of Fe, Mg, and P available.

#### 2.4.1 Point of zero charge, conductivity, and leachate

The BC, LDH/BC, and P-LDH/BC were tested to find their point of zero charge. The pH measurements were made using 0.1 mol L^-1^ NaCl aqueous solutions at pH 1, 3, 5, 7, 9, 11, and 13. The pH of the solution was adjusted with 1, 0.1, 0.01 mol L^-1^ NaOH and 1, 0.1, 0.01 mol L^-1^ HCl solutions. The initial and final pHs and electrical conductivities (EC) were measured in solutions with a Hanna pH/ORP meter. The solutions (25.0 mL) were mixed with 0.25 g of material in 2 replicates and stirred in an orbital shaker for 3 h. The supernatant was filtered, and the final pH and EC were recorded. The point of zero charge and conductivity was obtained by plotting the change in pH and EC, respectively, against the initial pH. The final solutions were analyzed on a PerkinElmer ELAN DRC II ICP-MS to determine the effect of pH on the amount of Fe, Mg, and P leached by the adsorbents.

#### 2.4.2 X-ray photoelectron spectroscopy

A Thermo Scientific K-Alpha XPS system was used for X-ray photoelectron spectroscopy (XPS) with a monochromatic X-ray source at 1486.6 eV, corresponding to the Al K11 line, with a spot size of 400 µm^2^. The constant analyzer mode was used to make the measurements, and the photoelectrons were collected at a takeoff angle of 900 relative to the overall sample’s fractal particle surface. The survey and high-resolution (HR) core level spectra were taken at a pass energy of 220 eV and 40 eV, respectively. The “Advantage v5.932” software provided with the instrument was used for XPS data acquisition.

#### 2.4.3 Surface area measurement

The surface areas and pore volumes of the BC, LDH/BC, and P-LDH/BC were measured by the N_2_ Brunauer-Emmett-Teller (BET) determinations. The BET was conducted using an N_2_ adsorption isotherm at ∼77 K (Micromeritics Tristar II Plus). Dubinin-Astakhov 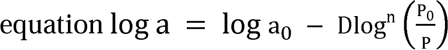, where ‘a’ denotes the amount of gas adsorbed per mass of adsorbent (mol/g), ‘a_0_’ is the micropore capacity (mol/g), ‘D’ is a constant, ‘P’ is the equilibrium pressure, and ‘P°’ denotes the saturation vapor pressure of adsorbate at temperature T (K). The surface area and pore volume of BC, LDH/BC, and P-LDH/BC are reported in Fig. S1.

#### 2.4.4 Ash and elemental composition

The carbon (C), hydrogen (H), nitrogen (N), and sulfur (S) elemental compositions of the BC, LDH/BC, and P-LDH/BC were measured by combustion analysis in the CHNS elemental analyzer. The ash content was determined by heating 0.5 g of the adsorbents to 750 °C for 2 h in air. The oxygen percentage of the adsorbents was determined by using Eq. 1.

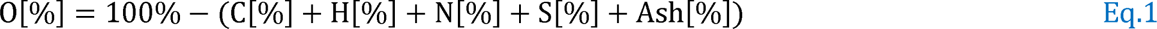

The metal contents in BC, LDH/BC, and P-LDH/BC were quantified by using PerkinElmer ELAN DRC II ICP-MS. Briefly, 0.10 g of the material was digested in HF, heated up to 150 °C with 25 mL of concentrated 3:1 (35% w/w HCl and 69% w/w HNO_3_) HCl: HNO_3_ mixture, diluted to 500 mL with 2% HNO_3_, and analyzed on the PerkinElmer ELAN DRC II ICP-MS.

### 2.5 Fixed-bed continuous flow column study

A fixed-bed continuous flow leaching experiment was performed to estimate the amount of P, Mg, and Fe leached by the absorbent. 2.0 g of P-Mg/Fe-LDH biochar was mixed with 10.0 g of soil and packed in a fixed-bed column (2.0 cm diameter x 7.0 cm length). The column was connected to a down-flow extraction setup with a flow rate of ∼1.5 mL min^-1^ (△P ∼ 1 mbar) at 25 °C using deionized water (∼pH 6.5). The leachate P, Mg, and Fe concentrations were measured using ICP-MS. Similarly, 0.169 g of TSP was mixed with 10.0 g of soil, and the P leaching rate was compared with P-Mg/Fe-LDH.

### 2.6 Greenhouse experimental setup

A Stough fine sandy loam (Coarse-loamy, siliceous, semiactive, thermic, Fragiaquic Paleudult) soil was collected from the Rodney Foil Plant Science Research Center at Mississippi State University. Topsoil was collected from 0-12 cm depth, hand-mixed to homogenize, air-dried for 72 h, and sieved through a 10 mesh sieve to remove any gravel and debris. The soil’s chemical properties were tested at the Mississippi State University Extension Service Soil Testing Laboratory to confirm a low to medium availability of extractable P level in the soil (Table S2) suitable for our experiment.

A greenhouse study was conducted at Mississippi State University between October 2022 and November 2022 to study the effects of P-LDH/BC on plant growth and nutrition. The study was conducted on common garden bush type beans (*Phaseolus vulgaris L.)*. ‘Blue Lake 156’ (bush type) variety was selected as the experimental crop. The treatments were arranged in a randomized complete block design, with 4.0 kg of soil in plastic pots with drainage. The amounts of fertilizer grade triple superphosphate (TSP), BC, LDH/BC, and P-LDH/BC are listed in Table 1.

**Table 1:**
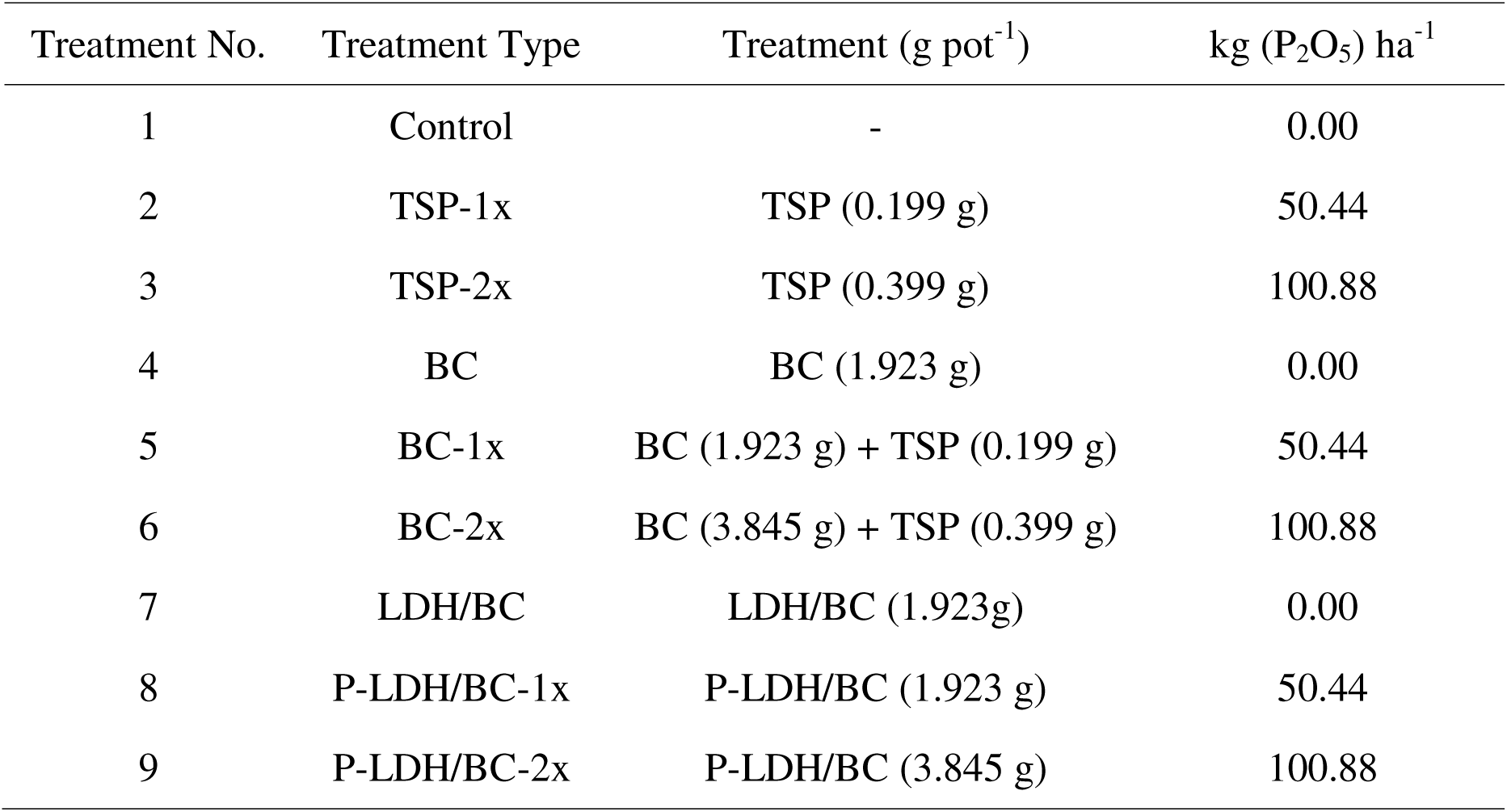
Treatments and P_2_O_5_ nutrient rate.

The treatments included 1) a control without fertilizer P or a biochar amendment; 2) control with fertilizer grade triple super phosphate (TSP), Ca(H_2_PO_4_)_2_ at a rate equivalent to 50.44 kg (P_2_O_5_) ha^-1^ (0.199 g per pot); 3) control with TSP as a P amendment at a rate equivalent to 100.88 kg (P_2_O_5_) ha^-1^ (0.399 g per pot); 4) untreated biochar without any P amendment (1.923 g BC per pot).;5) untreated biochar with TSP as a P amendment at a rate equivalent to 50.44 kg (P_2_O_5_) ha^-1^ (1.923 g biochar + 0.199 g TSP per pot); 6) untreated biochar with TSP as a P amendment at a rate equivalent to 100.88 kg (P_2_O_5_) ha^-1^ (3.845 g biochar + 0.399 g TSP per pot); 7) Mg/Fe-LDH biochar without any P amendment (1.923 g Mg/Fe-LDH biochar per pot); 8) P-Mg/Fe-LDH biochar as a P amendment at a rate equivalent to 50.44 kg (P_2_O_5_) ha^-1^ (1.923 g P-Mg/Fe-LDH biochar per pot);and 9) P-Mg/Fe-LDH biochar as a P amendment at a rate equivalent to 100.88 kg (P_2_O_5_) ha^-1^ (3.845 g P-Mg/Fe-LDH biochar per pot). All treatments were amended with N and potassium(K) fertilizer in the form of ammonium nitrate (35-0-0) and potassium chloride (0-0-60) at a rate equivalent to 56.04 kg (N) ha^-1^ (0.29 g NH_4_NO_3_ per pot) and 89.67 kg (K_2_O) ha^-1^ (0.272 g KCl per pot), respectively. To minimize confounding effects due to nutrient limitations, a blanked application of macro (except P) and micro-nutrients were applied to each treament. They were amended with Mg, lime, and micronutrients in the form of magnesium chloride, calcium carbonate, and Frit 503 (Frit Industries, Ozark Alabama) at a rate equivalent to 20.0 kg (Mg) ha^-1^ (0.31 g MgCl_2_ per pot), 1120.0 kg (CaCO_3_) ha^-1^ (0.31 g CaCO_3_ per pot), and 28.02 kg ha^-1^ (0.0509 g Frit 503 per pot). The pots were watered with DI water to achieve a moisture level of 65% field capacity, which was maintained by watering every 24 h a determined by weight. The pots were planted with 5 green bean seed, and the plants were thinned on day 7, leaving 2 healthy plants per pot. Each pot was innoculated with a rhizobial inncoulant mix consisiting of *Rhizobium leguminosarum* biovar. *viceae* (2×10^8^ cfu/g), *Bradyrhizobium* spp. (*vigna*) (2×10^8^ cfu/g), and *Rhizobium leguminosarum* biovar. *phaseoli* (2×10^8^ cfu/g) at the time of sowing. Additionally, nodules were observed in most plants during harvest sggesting successful innoculationa and engagement in nitrogen fixation. The Chlorophyll index (SPAD 503) and the plant height were measured on day 14, day 21, day 28, day 35, and day 42. Th bean pods were harvested on day 42, and their fresh weight was recorded immediately. The remaining plant parts were harvested on the same day by cutting them just above the soil level. The bean pods and the remaining plants were oven-dried separately at 65 °C for 72 h, and their dry weights were recorded.

### 2.7 Plant and soil nutrient analysis

The plants were separated into fruiting (bean pods) and vegetative or non-fruiting (stem, leaves, and flowers) tissue at harvest and analyzed separately. The tissues were oven-dried at 65 °C for 24 h, ground, passed through a 40-mesh sieve, and then re-dried at 65 °C for 24 h. The ground tissue was analyzed using a modified dry ashing procedure (Jones, 2001). The nutrient concentrations per gram of fruiting (bean pod) tissue and non-fruiting tissue were analyzed for macronutrients: P, Mg, Ca, K and micronutrients: Zn, Fe, B, Mn, Cu. Each nutrient concentration was multiplied by the respective tissue dry weight, and bean and plant results summed to obtain total mg of the nutrient uptake in mg per pot for each treatment. For example, Eq.2 shows calculation total P uptake per pot:

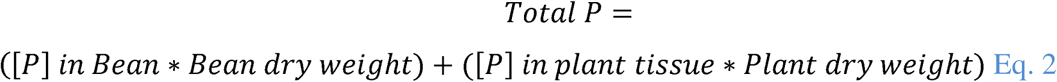

The P uptake efficiency for each treatment was calculated with Eq.3 as follows:

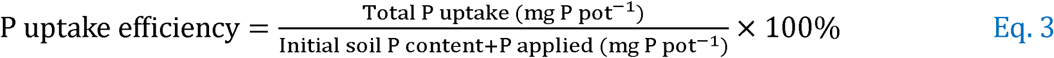

The initial air-dried soil nutrient content is mentioned in Table S2. The residual soil from each pot was air-dried after harvest and tested at the Mississippi State University Extension Service soil testing laboratory to quantify residual nutrients and determine the final soil pH.

### 2.8 Statistical analysis: Linear mixed models

The recorded data were analyzed in RStudio using R statistical software. Linear mixed-effect models using the “lme4” and “lmerTest” packages were employed to analyze variance (ANOVA) on the data. The ANOVA tested whether the administration of P through adsorbents (TSP, BC + TSP, or P-LDH/BC), the rate of P administered (0, 50.44, or 100.88 kg (P_2_O_5_) ha^-1^), or their interaction significantly affected all dependent variables of green beans and residual soil chemical properties. The P rates and the administration method were treated as fixed effects, while replicates were treated as random effects. The “Emmeans” package estimated and separated treatment means at p ≤ 0.05. All the statistical analyses were graphed using the “ggplot2” package.

## 3. Results and discussion

### 3.1 Characteristics of BC, LDH/BC, and P-LDH/BC

The C, H, O, N, and S mass percentages of dried BC were 84.74±2.76, 1.74±0.16, 10.96, 0.12±0.01, and 0.12±0.08, respectively (Table 2). The carbon percentages decreased to 46.68±4.95 and 46.86±3.50 during surface modification of the layered double hydroxide form and phosphate loading, respectively. The ash contents of BC, LDH/BC, and P-LDH/BC were analyzed using combustion analysis, as mentioned in Section 2.4.3. The ash percentages were subsequently increased from 2.32±0.28 to 24.94±0.31 after the modification of Mg/Fe layered double hydroxide on the BC surface. The subsequent ash percentage increment was 27.06±0.25 due to phosphate loading in P-LDH/BC. The BET surface areas of BC, LDH/BC, and P-LDH/BC were 603.9, 255.1, and 348.3 m^2^ g^-1^, respectively. The pore volumes were 0.042, 0.018, and 0.029 cm^3^ g^-1^ for BC, LDH/BC, and P-LDH/BC, respectively. The N%, P_2_O_5_%, and K_2_O% percentages of P-LDH/BC were 0.28, 4.77, and 0.19, respectively.

**Table 2.**
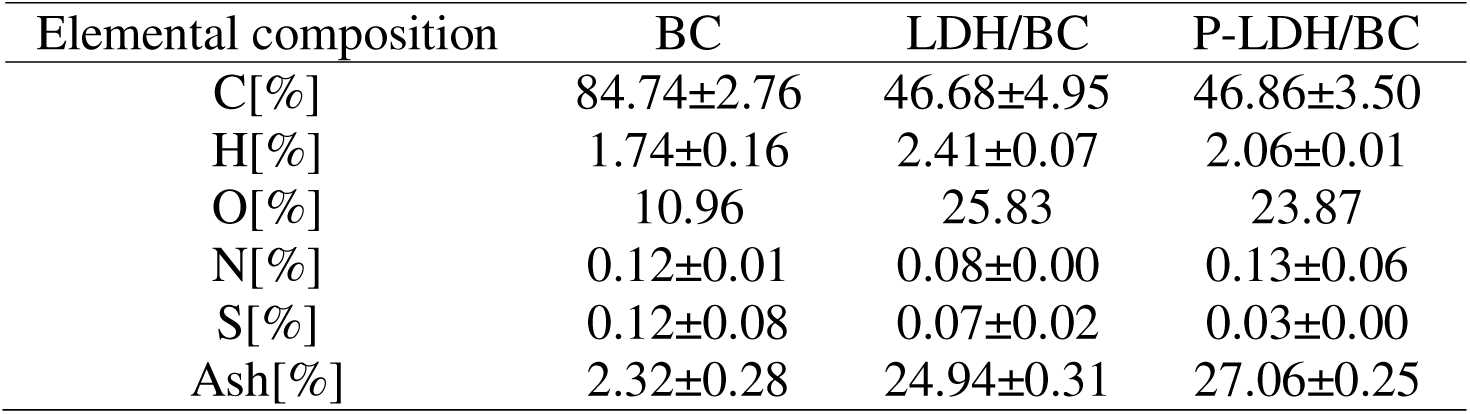
Elemental composition of BC, LDH/BC, and P-LDH/BC.

| Elemental composition | BC | LDH/BC | P-LDH/BC |
| --- | --- | --- | --- |
| C[%] | 84.74±2.76 | 46.68±4.95 | 46.86±3.50 |
| H[%] | 1.74±0.16 | 2.41±0.07 | 2.06±0.01 |
| O[%] | 10.96 | 25.83 | 23.87 |
| N[%] | 0.12±0.01 | 0.08±0.00 | 0.13±0.06 |
| S[%] | 0.12±0.08 | 0.07±0.02 | 0.03±0.00 |
| Ash[%] | 2.32±0.28 | 24.94±0.31 | 27.06±0.25 |

### 3.2 High-resolution X-ray photoelectron spectroscopy

The HR-XPS of Mg1s, Fe2p, O1s, and P2p provided the successful synthesis of Mg/Fe LDH on the biochar surface and the changes after phosphate adsorption. The HR Mg1s exhibited a clear progression of magnesium associations as the material changed from biochar to Mg/Fe LDH on biochar, then to phosphate-adsorbed Mg/Fe LDH. In pristine biochar, trace amounts of Mg were present with two small Mg1s components at 1303.76 eV (0.10%) and 1304.78 eV (0.09%), suggesting minimal, possibly surface-adsorbed Mg species on the biochar surface. After Mg/Fe LDH synthesis, a dominant Mg1s signal appeared at 1303.48 eV (7.03%) with a secondary component at 1304.28 eV (3.12%), consistent with Mg incorporated into brucite-like layers and environments typical of layered double hydroxides interacting with the carbonaceous surface. After the phosphate adsorption, the Mg1s features remained, with slight energy shifts and similar abundance (1304.28 eV at 7.90% and 1305.16 eV at 3.15%), suggesting that phosphate binding does not significantly displace Mg from its LDH environments but may slightly alter the surface Mg–O bonding, potentially through Mg–O–P interactions at the interface. Overall, the data showed a logical progression from trace Mg in biochar to significant Mg incorporation into Mg/Fe LDH, with only minor changes in the Mg environment upon phosphate uptake. This supports a phosphate-immobilization mechanism based on the LDH framework that maintains the Mg-hosting lattice.

The HR Fe2p spectra of Mg/Fe LDH on biochar indicated a mixed iron environment. Before phosphate exposure, the Fe2p spectrum showed Fe^2+^ and Fe^3+^ in the layered brucite-like structure, as well as surface oxide/hydroxide species bound to the biochar interface. After phosphate adsorption, the overall multiplet pattern remains, but a few components shift and change in intensity, signaling that the Fe–O bonds were perturbed by phosphate without changing the LDH framework, where the phosphate likely adsorbed onto Fe-oxide/hydroxide holes and possibly forms Fe–O–P interactions at the surface. These observations supported the successful synthesis of Mg/Fe LDH on biochar and revealed a phosphate-driven modification of the iron environment that aligns with known LDH-based phosphate immobilization mechanisms, without major lattice disruption.

HR-O1s spectra of pristine biochar, Mg/Fe LDH on biochar, and phosphate-adsorbed Mg/Fe LDH. In pristine biochar, three components appear: a lower-energy lattice/carbonate-associated O at 531.49 eV (5.65%), a mid-energy surface/moisture-related O at 533.21 eV (3.16%), and a higher-energy, weaker component around 535.71 eV (0.52%), collectively indicating a mixed oxygen landscape typical of carbonaceous materials with some surface functionalities. Upon synthesis of Mg/Fe LDH, the O1s distribution shifts markedly, with a dominant Brønsted/Lewis-oxide signature at 531.65 eV (35.18%) and a residual lattice-like component at 529.57 eV (1.56%), reflecting the incorporation of brucite-like hydroxide layers and new Mg/Fe–O environments interfacing with the carbon substrate. After phosphate adsorption, the spectrum remains dominated by the 531.71 eV component (38.71%), with a secondary contribution at 533.09 eV (10.01%), consistent with additional phosphate-related oxygen species (P–O–Mg/Fe linkages or adsorbed phosphate) coexisting with the continued LDH-derived oxygen environments. These Mg/Fe LDHs synthesized on biochar were successful and indicate phosphate immobilization mechanisms involving Mg/Fe–O–P interactions while preserving the LDH-typical oxygen framework.

The P2p signal after phosphate adsorption on Mg/Fe LDH with biochar was characterized by a distinct P2p3/2 component at 133.78 eV (8.84%), consistent with phosphate in a bridging environment bound to metal oxygens. This peak, together with the absence of a contrasting higher-energy P2p, supports the dominance of surface-bound phosphate species engaged in inner-sphere interactions with Mg–O and Fe–O sites, likely forming P–O–Mg/Fe linkages at the LDH/biochar interface. The appearance of a pronounced P signal at this binding energy, alongside unchanged shifts in the Mg, Fe, and O signals, indicates successful phosphate immobilization without disrupting the LDH lattice, consistent with a mechanism in which phosphate is anchored at surface oxygens within or atop the brucite-like layers. Overall, effective phosphate incorporation into the LDH framework contributes to immobilization on the Mg/Fe LDH-biochar composite.

### 3.3 Effect of pH

The changes in pH (ΔpH = final pH – initial pH) and conductivity (ΔConductivity = final conductivity – initial conductivity) from pH 1 to 13 were measured in 0.01 mol L^-1^ NaCl solution matrices (Fig. 3[A] and Fig. 3[B]). Soil, BC, LDH/BC, and P-LDH/BC showed PZC of 7.28, 8.95, 10.69, and 10.32, respectively (Fig. 3[A]). The PZC of BC is attributed to the presence of carbonates and oxides of Mg, Ca, and K formed during pyrolysis, which are hydrolyzed upon contact with water (Rodrigo et al., 2024). The LDH/BC matric has increased PZC due to the fundamental nature of the LDH structure, which is directed by the presence of OH^-^ ions. P-LDH/BC exhibits a decrease in PZC due to the introduction of additional negative surface charges by HPO_4_^2-^. The pH changes of soil, BC, LDH/BC, and P-LDH/BC were 0.04, 0.08, 6.71, and 5.57 at initial pH values of 1.44, 1.5, 1.5, and 1.5, respectively. LDH/BC and P-LDH/BC showed increased pH changes due to the higher amounts of H^+^ neutralized in 0.1 mol L^-1^ NaCl medium at pH 1.5 due to the release of OH^-^ by the LDH structure. However, the P-LDH/BC showed a lower change in pH due to the reduced capacity to release OH^-^ ions after phosphate adsorption on the LDH/BC.

**Fig. 1.**
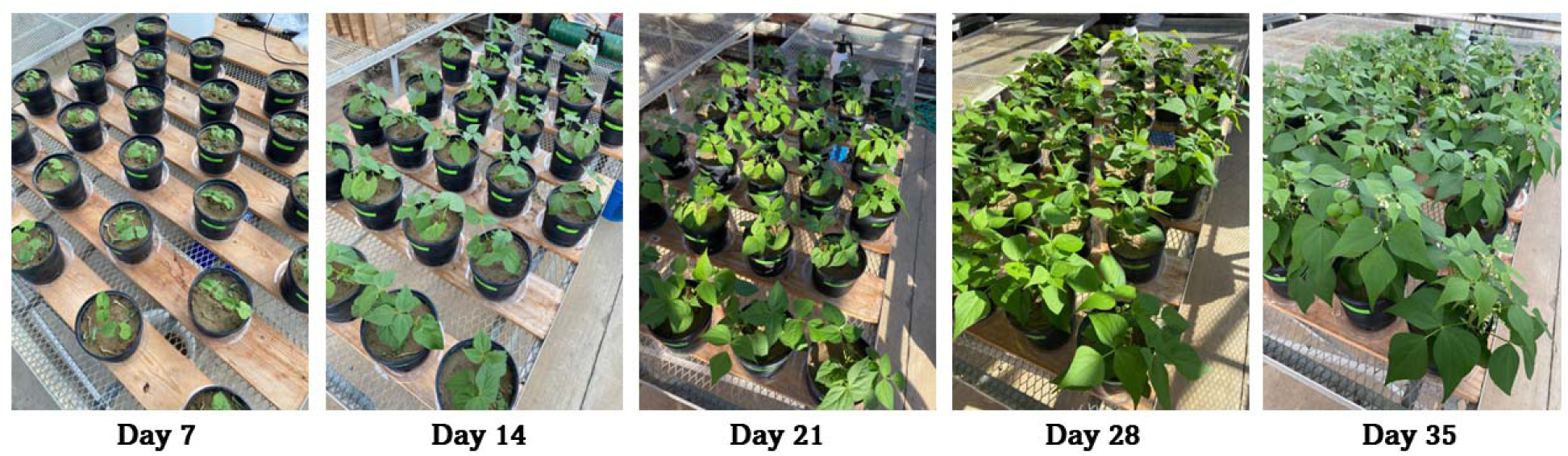
Green Beans (*Phaseolus vulgaris L.)* plants after Days 7, 14, 21, 28, and 35

**Fig. 2.**
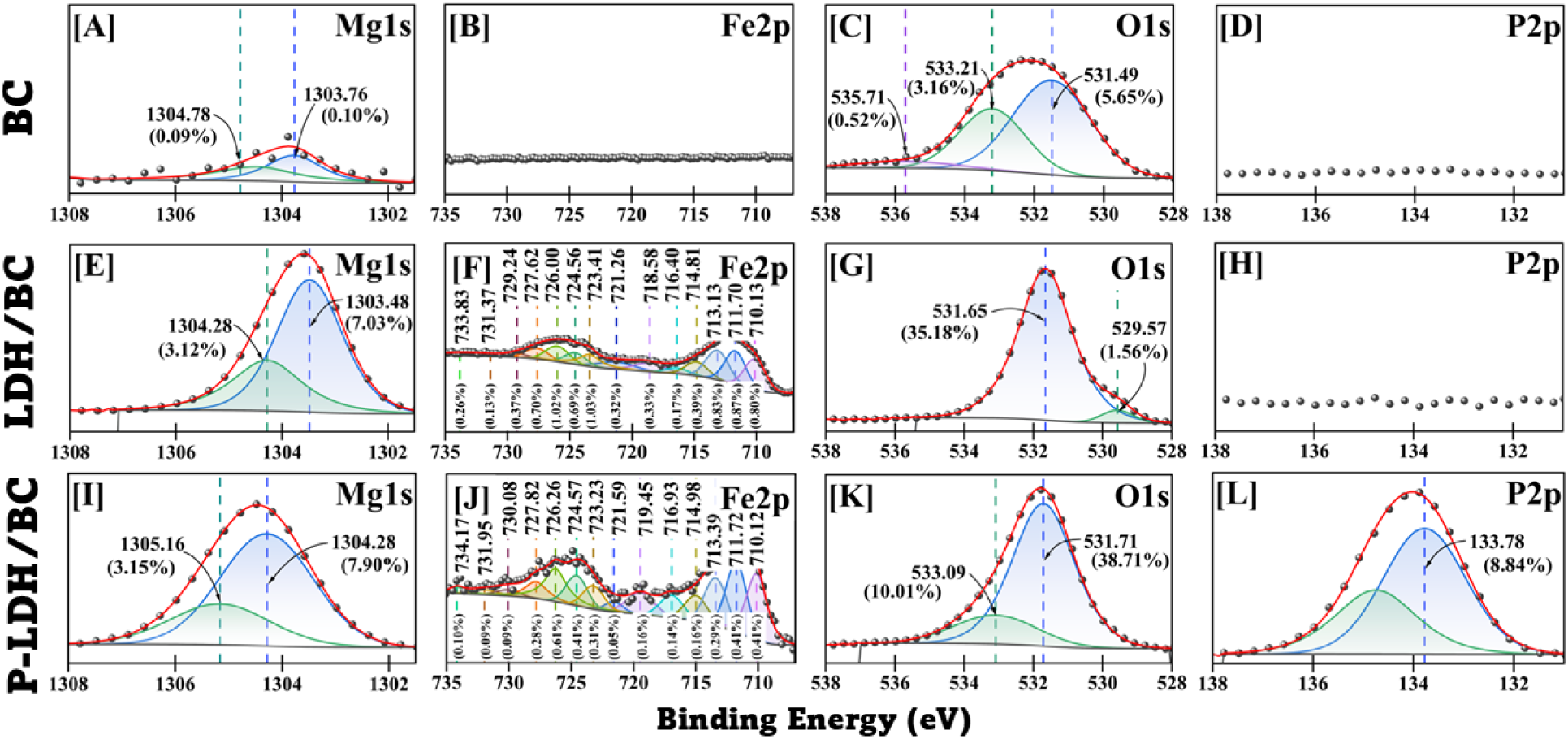
High-resolution Mg1s, Fe2p, O1s, and P2p XPS spectra for (A)-(D) BC, (E)-(H) LDH/BC, and (I)-(L) P-LDH/BC, respectively.

**Fig. 3.**
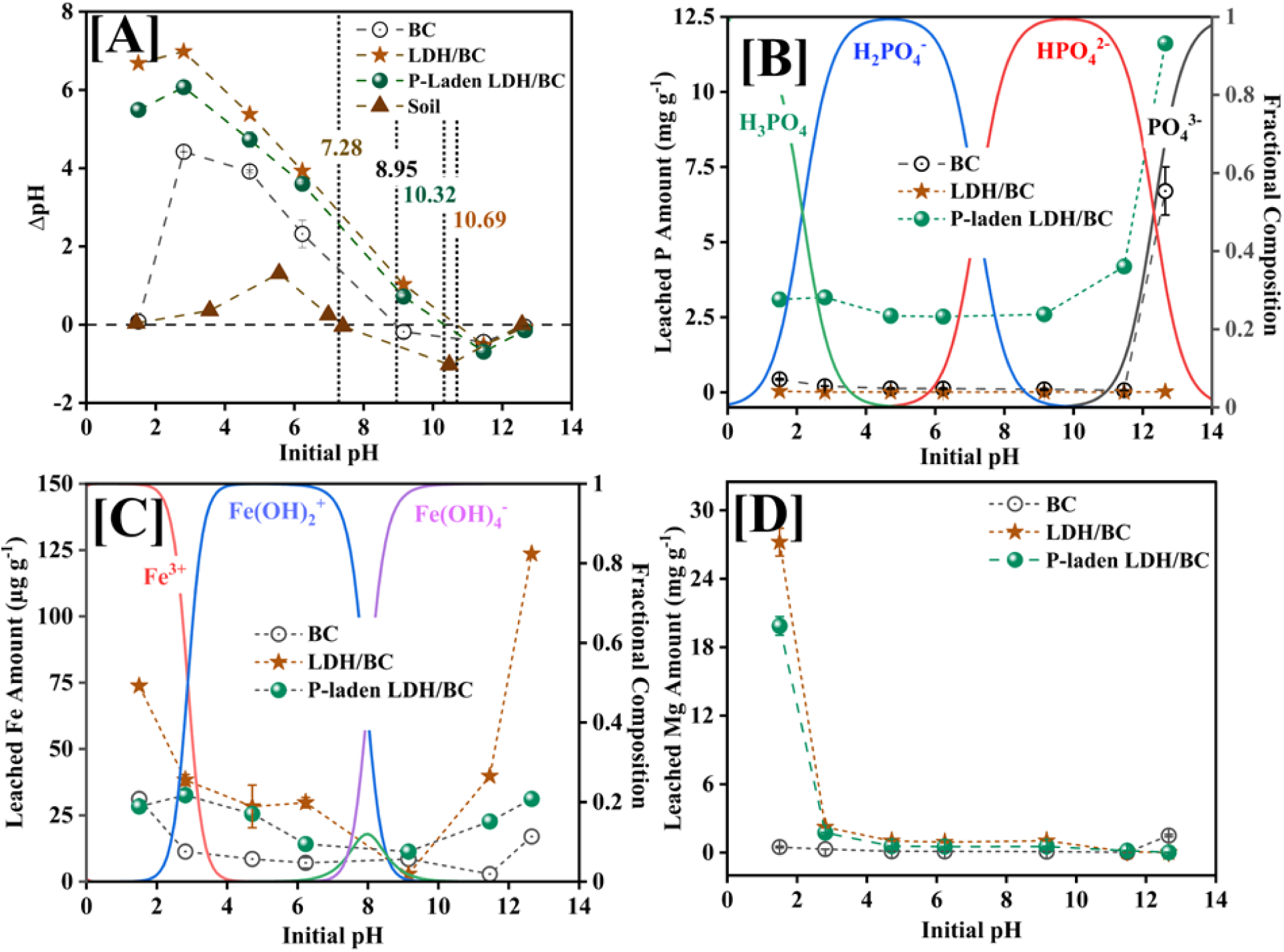
[A] *The pH change,* [B] *P,* [C] *Fe, and* [D] *Mg leaching versus initial solution pH for BC, LDH/BC, and P-laden LDH/BC*

The Δconductivities of BC, LDH/BC, and P-LDH/BC were −8.0, −13.55, and −13.9 mS cm^-1^, respectively, at an initial pH of 1.5. The LDH/BC and P-LDH/BC exhibited larger negative conductivity changes at an initial pH of 1.5, due to the neutralization of protons by OH- released and the adsorption of Cl^-^ ions, indicative of their high anion exchange capacities. The release of 3090 μg g^-1^ of PO_4_^3-^ ions at pH 1.5 (Fig. 3[C]) by P-LDH/BC directed the pH change by buffering (H_2_PO_4_^-^/H_3_PO_4_). Additionally, the adsorption of Cl^-^ significantly reduced the conductivity of the medium.

BC, LDH/BC, and P-LDH/BC exhibited pH changes of −0.05, −0.07, and −0.14, respectively, at an initial pH of 12.65. The Δconductivities of BC, LDH/BC, and P-LDH/BC were −5.89, −5.71, and −7.06 mS cm^-1^, respectively, at an initial pH of 12.65. The anion exchange capacity increased in LDH/BC ≈ BC < P-LDH/BC at a high pH of 12.65. The observed changes in conductivity indicate that the anion exchange capacity increased in the order of BC << LDH/BC < P-LDH/BC at a low pH of 1.5. P-LDH/BC had the highest anion exchange capacity, enhanced by pre-adsorbed phosphate leaching (Fig. 3[C]).

The PO_4_^3-^, Fe^3+^, and Mg^2+^ ion leaching amounts were measured from an initial pH of 1.5 to 12.65 (Fig. 3[C]-[E]). The PO_4_^3-^, Fe^3+^, and Mg^2+^ leached at 3090, 28, and 19868 μg g^-1^ of P-LDH/BC at an initial pH of 1.5. The PO_4_^3-^ leached at a significantly lower rate of 433 μg g^-1^ of BC and was the weakest at 36 μg g^-1^ of LDH/BC at an initial pH of 1.5 due to the phosphate retention by the positive surface charges of LDH/BC. The Fe^3+^ and Mg^2+^ leached at a higher rate of 74 and 27210 μg g^-1^ of LDH/BC, respectively, accompanied by an OH^-^ release at a low pH of 1.5, disrupting the structural integrity of the layered double hydroxide. The Fe^3+^ and Mg^2+^ leached at lower rates from P-LDH/BC at pH 1.5 due to the stabilizing effect of pre-adsorbed phosphate. The BC leached Fe^3+^ and Mg^2+^ at concentrations of 31 and 492 μg g^-1^, respectively, at an initial pH of 1.5.

### 3.4 Fixed-bed continuous flow study

Fixed-bed continuous-flow treatment was used to study the leaching behavior. A 1:5 mass ratio of P-LDH/BC to soil was used in the leaching study, with a P loading of 41.7 mg relative to the total weight. The P leaching rate of P-LDH/BC was compared with TSP (41.7 mg of P loading) at a ∼1.5 mL min^-1^ flow rate (Fig. S3). TSP exhibited a higher P leaching rate than P-LDH/BC. A ∼19.3 mg of P was leached from TSP at 2 L of leachate, which was 1.4 time higher P leaching compared to P-LDH/BC. The slopes of the leaching curve depend on the flow rate, nutrient concentrations (PO_4_^3-^, Mg^2+^, and Fe^3+^), mass transfer rate, pore diffusion rates, and soil exchange rate. For P-LDH/BC, the cumulative leached P amounts were 7.8, 11.2, 13.0, and 14.0 mg, respectively, after passing 0.5, 1, 1.5, and 2.0 L (Fig. 4(A)). The P-leached percentages were ∼19, ∼27, ∼31, and ∼34%, respectively. The higher P leaching rate was observed until passing 1.0 L due to the loosely bound H_2_PO_4_^-^ and HPO_4_^2-^ on the LDH/BC surface. The Mg leaching amounts were 0.11, 0.20, 0.27, and 0.31 mmol after 0.5, 1, 1.5, and 2.0 L passed, respectively (Fig. 4(B)). However, Fe leaching was significantly lower in pH 6.5 DI water, approximately 238 times lower than Mg leaching after 2.0 L passed.

**Fig. 4.**
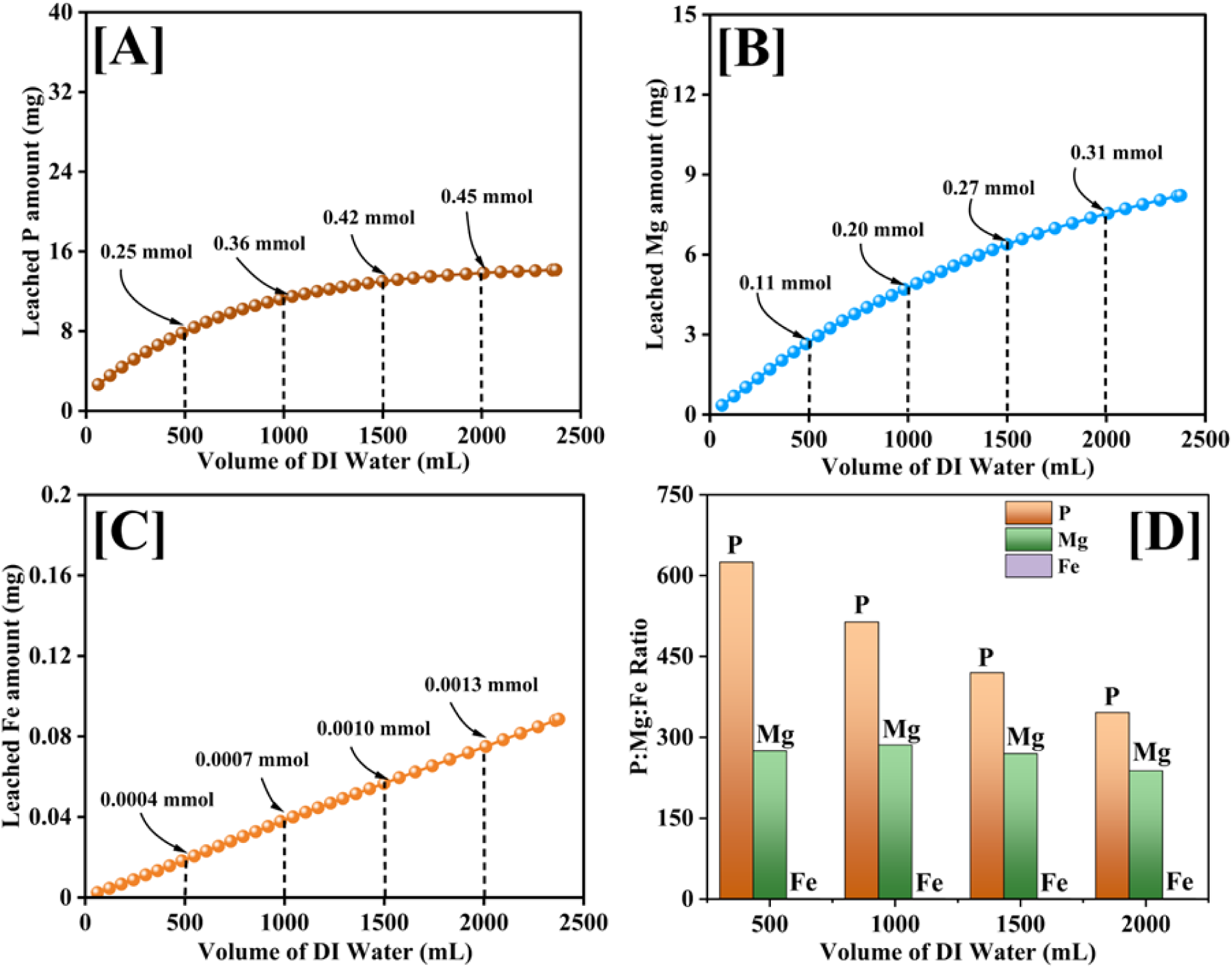
[A] *P,* [B] *Mg,* [C] *Fe leaching amounts over DI water passed through the column*, [D] *P:Mg:Fe leached ratio after 500, 1000, 1500, and 2000 mL of DI water passed*.

### 3.5 Plant height and chlorophyll content

The effects of treatments on plant height is shown at weekly intervals, starting with 14 days after planting in Fig. 5 (A). The standard deviation in average plant height for all treatments up to 21 days (1.3 cm) was relatively minor compared to differences observed after 35 (3.8 cm) and 42 (3.1 cm) days of growth, indicating that the treatment best predicted variance in plant height. The minor initial deviation was likely due to adequate availability of soil nutrient realtive to plant demand during early growth stages. The average height of plants treated with P-LDH/BC at a P_2_O_5_ rate of 100.88 kg ha^-1^ was significantly taller than other treatments after day 35 (32.8 cm, p = 7.088e-03) and 42 (35.5 cm, p = 6.758e-03). The continued growth may be due to the controlled release of P and other nutrients from P-LDH/BC. The significantly (p = 1.258e-10) higher levels of Mg in the residual soil in treatments with P-LDH/BC (Table S4) suggests additional Mg leaching from the LDH which could also result in the increased growth in treatment 9. Overall, plant heights increased with higher rates of P across TSP, BC + TSP, and P-LDH/BC treatments after days 28 (p = 3.95e-04), 35 (p = 5.969e-05), and 42 (p = 8.14e-04), due to the greater amounts of P applied indicating a positive response to applied P for this soil.

**Fig. 5.**
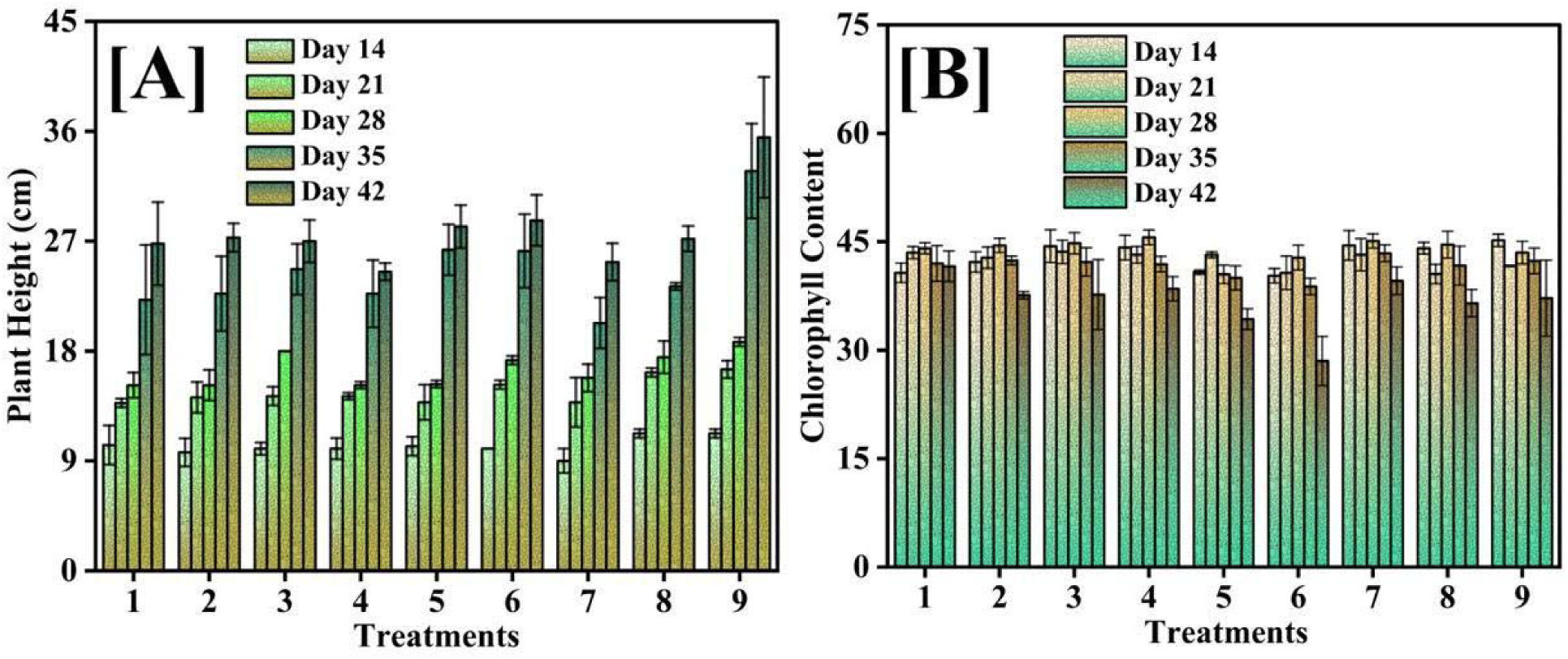
[A] Plant height and [B] chlorophyll content after days 14, 21, 28, 35, and 42.

Higher chlorophyll content indicates carbohydrate production during photosynthesis and overall plant health, with magnesium essential for photosynthetic activity and chlorophyll synthesis (Ahmed et al., 2020; Farhat et al., 2016). Magnesium is highly mobile in plants and crucial for the source-to-sink transport of carbohydrates (Farhat et al., 2016). This study applied magnesium to all the pots before sowing as MgCl_2_ at a rate of 20 kg ha^-1^. Chlorophyll readings were taken weekly, starting with 14 days after sowing, from the middle part of the leaf (Fig. 5[B]). The chlorophyll concentrations declined across all treatments between days 28 and 42 but the magnitude of decline was less pronounced in treatments without phosphorus amendments, across the different treatment types (TSP, BC + TSP, and LDH/BC; Fig. 5[B]). The chlorophyll concentrations were significantly higher in plants treated with lower P-rates across the different treatement types (TSP, BC + TSP, and LDH/BC) after 14 (p = 2.028e-02), 28 (p = 3.747e-02), and 42 days (p = 5.086e-03, Fig. S4[A]). The higher concentrations can be attributed to the lack of Pi-dependent metabolic functions, particularly Rubisco and fructose-1, 6-bisphosphatase, under low P stress to maintain photosynthesis (Bechtaoui et al., 2021). In a 2021 review, Bechtaoui et al. reported total chlorophyll content in leguminous plants was higher in P-deficient plants than those with sufficient P, with a reduction of −46% and −31% for soybean and cowpea, respectively (Bechtaoui et al., 2021). Also, low P is known to cause stunted growth, shorter leaves, and dark green foliage (Malhotra et al., 2018), potentially increasing chlorophyll content per unit leaf area. Despite increased chlorophyll concentrations, plants without phosphorus amendment accumulated significantly lower biomass, and thus the chlorophyll measurement was not directly related to plant growth and biomass acculmulation, as indicated by plant dry weight (p = 4.983e-07, Fig. S4[B]) and bean dry weight measurements (p = 1.46e-06, Fig. S4[C]).

### 3.6 Plant dry weight, bean fresh weight, bean dry weight, and soil pH

Bean pods with developing seed were separated from the rest of the plant vegetative structures at harvest, and both were dried separately at 65 °C for 72 h. The average plant dry weight, bean fresh weight, and bean dry weight for treatments 1 to 7 are reported in Fig. 6[A-C]. A significant relationship was observed between yield and P amendment rates, where plant dry weight (p = 4.983e-07), bean fresh weight (p = 3.126e-06), and bean dry weight (p = 1.46e-06) increased with increasing rates of P across the different treatment types (TSP, BC + TSP, and P-LDH/BC).

**Fig. 6:**
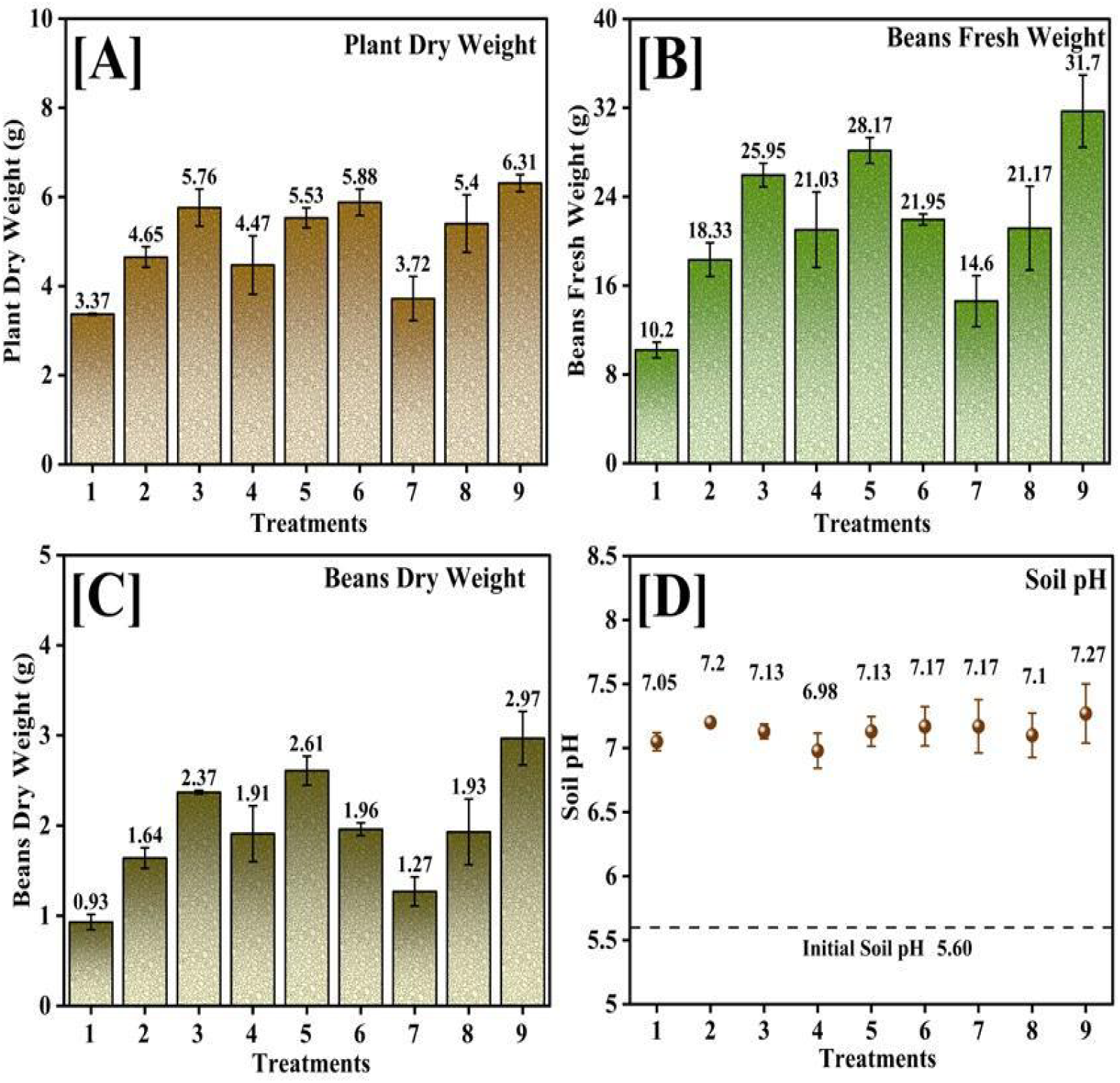
[A] *Plant dry weight*, [B] *Beans fresh weight*, [C] *Beans dry weight*, *and* [D] *residual soil pH*

The plants treated with BC and P-LDH/BC had increased yields of plant dry weight (p = 1.056e-02), bean fresh weight (p = 6.046e-04), and bean dry weight (p = 1.788e-04) at P_2_O_5_ rates of 0 and 50.44 kg ha^-1^. BC’s nutrient retention and controlled-release effects are conssitent with the increased yields. The untreated BC has a greater surface area available to retain highly soluble TSP and to reduce the solubility of added nutrients (K^+^, Mg^2+^, Ca^2+^, and micronutrients), making them available for growth at later stages. Plants treated with P-LDH/BC at P_2_O_5_ rates of 100.88 kg ha^-1^ had the highest overall measurements for average plant dry weight (6.31 g, p = 1.056e-02), bean fresh weight (31.7 g, 6.046e-04), and bean dry weight (2.97 g, p = 1.788e-04). We speculate that a controlled release of adsorbed P from P-LDH/BC provided continuous support for plant growth and improved yields. Overall, the increased height and weight yields with higher P amendment rates reinforce the key contribution of P to plant growth. The highest yields for all height and weight parameters were recorded for plants treated with P-LDH/BC at a P_2_O_5_ rate of 100.88 kg ha^-1^, demonstrating the benefits of the controlled release properties of P-LDH/BC for sustained growth and crop yields.

Soil pH changes depend on a variety of biological, chemical, and physical factors. The soil’s initial pH was 5.60, and the final pH values for all treatments became basic during the greenhouse study period, but no significant differences were observed between treatments Fig. 6(D). Treatment 9 had the slightly highest pH due to the combined effect of a higher biochar application rate and the highest HPO ^2-^ ammendment.

### 3.7 Macronutrient uptake: P, Mg, K, and Ca

Bean pods were separated from the vegetative plant tissue (non-fruiting) at harvest and processed separately to evaluate commercial yield and nutritional contents. The average P content for the bean pods is reported in Fig. 7[C]; for the remaining plant tissue see Fig. 7[A]. The P content increased for both bean pods (p = 6.825e-04) and the remaining plant tissue (p = 2.125e-04) as P amendment rates increased across the different treatment types (TSP, BC + TSP, P-LDH/BC; Fig. S5[A]). Plants treated with P-LDH/BC had a higher P content for the beans (p = 5.353e-03) due to its controlled-release capacity. The total P uptake calculated using Eq. 2 (see section 2.7 *Plant and soil nutrient analysis*) for treatments 1 to 9 was 7.95, 11.70, 17.75, 9.57, 15.56, 17.59, 8.53, 14.37, and 20.94 mg (P) pot^-1^, respectively; thus P-LDH/BC had the highest P content (p = 5.703e-02, sub-significant) at a P_2_O_5_ rate of 100.88 kg ha^-1^.

**Fig. 7:**
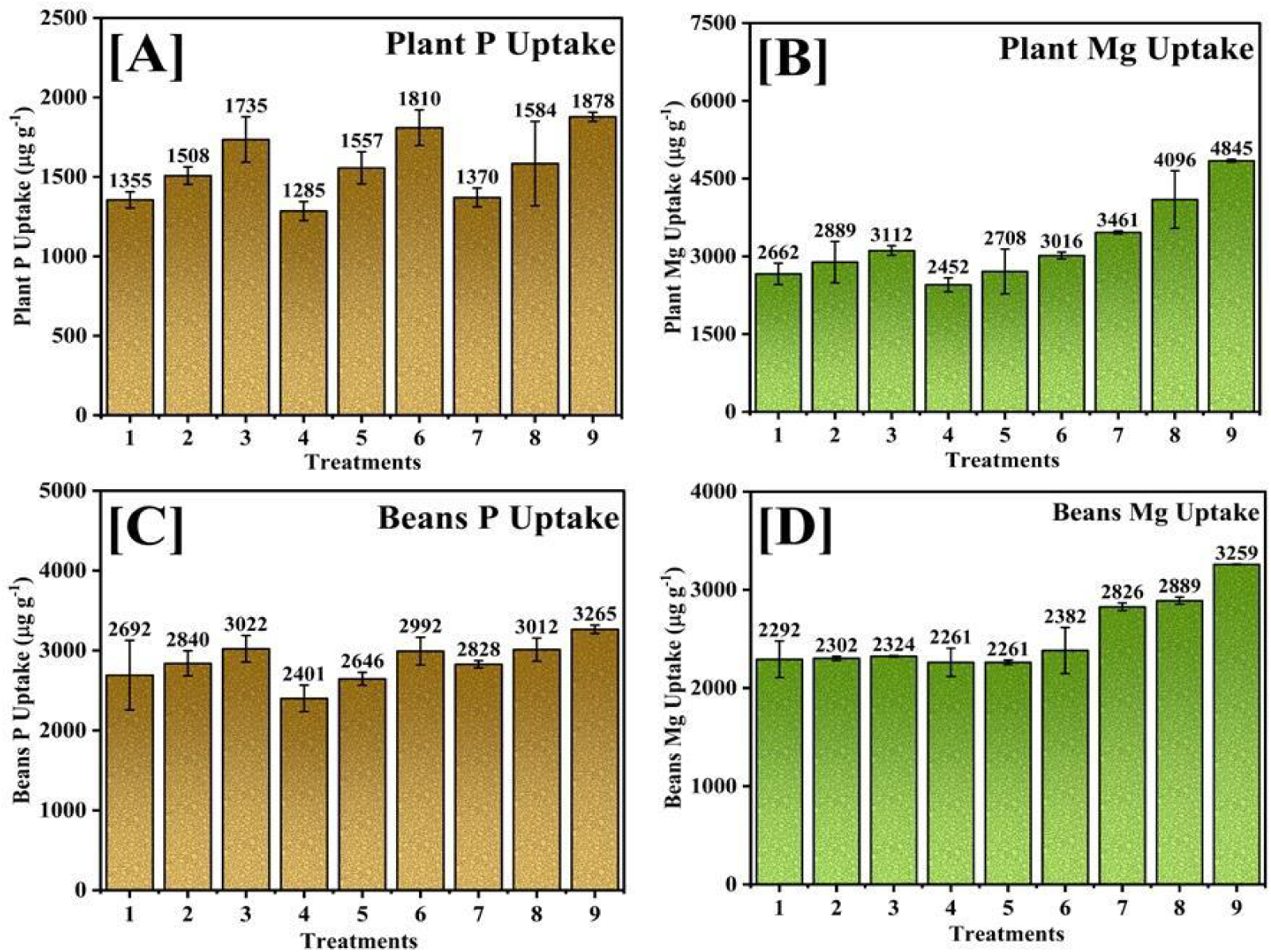
*Plant* [A] *P and* [B] *Mg uptake, Beans* [C] *P and* [D] *Mg uptake from 1-9 treatments*

The P uptake efficiency was calculated with Eq. 3 (see section 2.7 *Plant and soil nutrient analysis*) to normalize against total P present per pot and measure differences across different treatment types (TSP, BC + TSP, P-LDH/BC). The P uptake efficiency for treatment 1-9 were 4.16%, 4.21%, 4.86%, 5.01%, 5.60%, 4.82%, 4.46%, 5.17%, and 5.74%, respectively with no significant difference (p = 0.29) observed across different treatment types (TSP, BC + TSP, P-LDH/BC). Several factors, including soil pH, temperature, microbial composition, microbial symbiosis, and plant physiological processes, impact P use efficiency in plants (Miyasaka et al., 2001; van de Wiel et al., 2016; Veneklaas et al., 2012). The plants treated with LDH/BC composites without adsorbed P had a lower P uptake efficiency (4.46%) than those treated with BC without additional P. The adverse effects were expected due to the capacity of LDH/BC to exchange and retain soluble P from the soil (Navarathna et al., 2022). The P uptake efficiency increased as the P-LDH/BC amendment increased, thus treatment 9 showed the highest efficiency (5.74%), suggesting improved labile P availability and increased microbial activity. The variability within treatments prevented a statistically significant conclusion, and analysis of microbial community structures might help explain the differences. Further investigation is required to draw definitive conclusions about the impact of different treatment types on P uptake efficiency.

The Mg content in residual soil varied significantly (p = 1.258e-10; Table S4) between different treatment types, prompting a deeper investigation of uptake effects in plants. The initial soil had a low to medium soil test value of 40.35 kg Mg ha^-1^ (Table S2). Additionally, MgCl_2_ was applied to all treatments at a rate of 20.0 kg Mg ha^-1^, equivalent to 78.1 mg of Mg per pot. The average Mg content for the bean pods from treatment 1 to 9 is reported in Fig 7[D]; for the remaining plant tissue see Fig. 7[B]. The plants treated with P-LDH/BC had a higher Mg content for the bean (p = 8.472e-06) and in the remaining plant tissue (p = 3.513e-06). The concentration of Mg increased with increasing P-LDH/BC amendment rates in the bean (p = 6.487e-02) and the remaining plant tissue (1.078e-03). The higher Mg content in plants treated with P-LDH/BC translates to improved seed quality, enhanced germination, and increased nutritional content for human consumption (Gerendás et al., 2013). Mg is one of the essential micronutrients lacking in human diets worldwide, posing the risk of critical deficiencies (Broadley et al., 2010; Rosanoff, 2013). A higher Mg content in edible plant tissue can help address hidden hunger and nutrient malnutrition in humans (White et al., 2009). Mg is vital in plant carbohydrate formation and protein biosynthesis, influencing the quality and nutritional value of various crops (Gerendás et al., 2013).

The total average Mg uptake for treatments 1 to 9 was 11.14, 16.45, 20.75, 14.01, 19.80, 23.11, 17.89, 29.26, and 43.38 mg (Mg) pot^-1^; thus plants treated with P-LDH/BC showed higher average Mg uptake (p = 4.529e-05), which increased with increasing P_2_O_5_ amendments (p = 1.419e-05). The total Mg uptake was influenced by an interaction between treatment and rate effects (p = 7.611e-03), with plants treated with P-LDH/BC at 100.88 kg (P_2_O_5_) ha^-1^ exhibiting the highest uptake (Fig. S5[B]).

Treatment 8 was amended with 1.923 g of P-LDH/BC at a rate of 50.44 kg (P_2_O_5_) ha^-1,^ while treatment 7 received 1.923 g of LDH/BC with no P_2_O_5_ amendments. Both treatments showed statistically similar Mg uptake, regardless of the P amendment applied to the soil. Treatment 9 was amended with 3.845g of P-LDH/BC at a rate of 100.88 kg (P_2_O_5_) ha^-1^ and had significantly higher Mg uptakes (Fig. S5[B]). This result established that the higher uptakes were due to additional Mg leached from the LDH structure. The residual soil Mg after harvest was significantly higher for treatments 7, 8, and 9 amended with P-LDH/BC (p = 1.258e-10), with the highest value for treatment 9 (p = 5.124e-04), which received double the amount of P-LDH/BC of the three. The residual Mg amounts were 92.2, 87.75, and 131.12 kg (Mg) ha^-1^, significantly higher than the total (initial + amended) amount of 60.35 kg (Mg) ha^-1^ in the pots. These results indicate that the increased Mg uptake was primarily due to additional Mg leaching from the LDH structure, as well as the possible improvement in soil conditions and microbial activity resulting from the BC component in the P-LDH/BC.

K and Ca levels in the bean pods and other vegetative plant tissue were analyzed to understand the effect of the synthesized P-LDH/BC fertilizer on comprehensive macronutrient uptake and tissue nutrient composition in green beans.

The starting soil had a K content of ∼80.7 kg ha^-1^ (146.69 mg (K) pot^-1^). K was applied as KCl to all treatments at a rate of 89.67 kg (K_2_O) ha^-1^, equivalent to 142.5 mg (K) pot^-1^. Average K uptake for bean pods and other vegetative tissue varied between 18.52 - 22.21 mg g^-1^ and 8.95 and 14.92 mg g^-1^of dried tissue, respectively for all treatments. A significant relationship was observed between K uptake and increasing P amendment rates. The total K uptake per pot increased (p = 1.171e-02) while the concentration of K per gram of tissue decreased in both the beans (p = 2.101e-03) and the remaining plant tissue (p = 2.562e-05) with an increasing rate of P amendment. The reduced K concentrations can be explained by a combination of increased magnesium levels in soil (Kasinath et al., 2014) and a dilution effect (Sandaña et al., 2024), where the enhanced growth and biomass accumulation reduced the concentration of K in the plant tissue. The ratio of K: Mg uptake per gram of dried tissue was calculated for both the beans and the remaining plant tissue. All treatments 1-9 were amended with a 3:1 ratio of K to Mg, as suggested by soil analysis results for green beans. The ratio of K: Mg concentrations per gram of dried tissue, however, decreased with increasing P amendment rates in the beans (p = 1.025e-02) and the remaining plant tissue (p = 3.840e-04). The plants treated with P-LDH/BC had significantly lower K: Mg ratios than other treatments in beans (p= 5.535e-04) and in the remaining plant tissue (p= 4.911e-03). These treatments, as shown above, had the highest biomass and Mg uptake due to Mg leached from the LDH component. K and Mg are cationic ions, and uptake can be competitive and antagonistic in some plants (Xie et al., 2021). The total K uptake per pot increased with increasing P amendment rates. However, higher accumulated biomass resulted in a lower K concentration per gram of dried tissue, confirming a strong dilution effect that may be influenced by increased Mg amendment.

The total average Ca uptake per pot increased with increasing P amendment rates (p = 7.166e-04) and did not differ significantly among TSP, BC + TSP, and P-LDH/BC treatments. The concentration of Ca per gram of dried tissue in the bean (p = 9.896e-06) and the remaining plant tissue (p = 4.529e-03) was higher for the plants treated with TSP. The plants treated with TSP at 50.44 kg (P_2_O_5_) ha^-1^ had the highest Ca uptake for the beans (p = 5.335e-04). The Ca concentrations in beans increased in the order of plants treated with P-LDH/BC < BC + TSP < TSP (p = 9.896e-06).

### 3.8 Micronutrient uptake: Fe, Zn, B, Mn, Cu

Micronutrient play an essential role in normal plant development and metobolic function. The Fe leaching from P-LDH/BC (shown above; Fig. 4[C]) prompted a comprehensive analysis of micronutrient levels in the plant tissue. Micronutrient uptake depends on fertilizer application rate, soil conditions, and plant types (Kochian, 1991). Frit is one of four major types of micronutrients (Mortvedt, 1991). Frit was applied at a rate of 28.02 kg ha^-1^. Fig. 8[A]-[E] summarizes the nutrient uptake of Zn, Fe, B, Mn, and Cu, respectively, for beans and the remaining plant tissue in μg g^-1^ of dried tissue. The rate of uptake for the micronutrients is mentioned in Table S5-6. In beans, the rate of Fe uptake increased with increased rate of P amendment across all treatment types: TSP, BC + TSP, and P-LDH/BC (p = 1.527e-4) (Table S6). The average rate of Fe uptake was highest at 78.88 μg (Fe) g^-1^ of dried tissue in beans harvested from plants treated with P-LDH/BC, treatment 9. The increment is due to increased iron availability leached from 3.845 g of P-LDH/BC applied to the treatment. Mn uptake rates increased with higher TSP and BC + TSP amendment rates but remained unchanged with higher P-LDH/BC amendment rates (p = 1.229e-2). The total average manganese uptake per pot increased with increasing P amendment rates (p = 2.324e-05) and did not differ significantly among TSP, BC + TSP, and P-LDH/BC treatments.

**Fig. 8.**
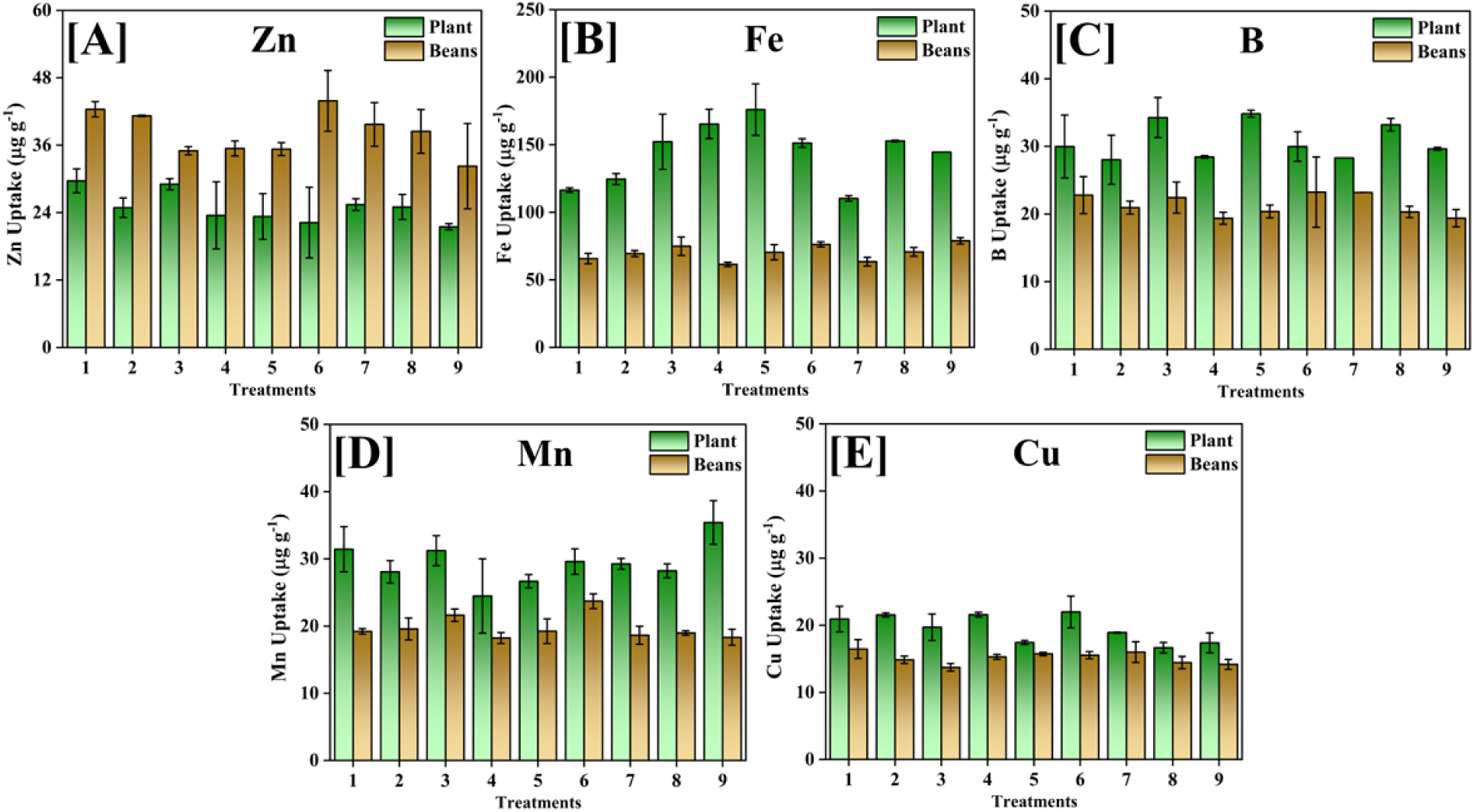
[A] *Zn*, [B] *Fe*, [C] *B*, [D] *Mn, and* [E] *Cu uptake from plants and beans from treatments 1-9*.

## 4. Conclusion

This work advances sustainable nutrient management by integrating phosphorus-loaded Mg/Fe layered double hydroxides (LDH) with Douglas fir biochar to create a novel, controlled-release fertilizer system. By directly synthesizing Mg/Fe-LDH on biochar via co-precipitation, loading the composite with phosphate through anion exchange, and characterizing the resulting material with elemental analysis, BET surface area measurements, and X-ray photoelectron spectroscopy, a material that combines high surface area, accessible active sites, and enhanced dispersibility was established. The resulting P-Mg/Fe-LDH biochar exhibited phosphate-buffering capacity and improved dispersibility compared with conventional LDHs, addressing a key limitation of LDH-based sorbents. This multifunctional material aligns with a broader shift toward precision fertilization, in which nutrient release is tuned to plant demand and environmental conditions, potentially reducing leaching losses that contribute to eutrophication and hypereutrophication in aquatic systems.

Greenhouse results with bush beans (*Phaseolus vulgaris L.*) demonstrate the practical benefits of this composite. Plants grown with P-Mg/Fe-LDH biochar showed measurable improvements in growth metrics, including increased yield, biomass, plant height, and phosphorus uptake, at application rates that reflect realistic agricultural practice. Notably, bean fresh weight, plant dry weight, plant height, and the phosphorus uptake were considerably higher, suggesting that the controlled-release mechanism maintains nutrient availability over critical growth stages while limiting excess phosphorus entering the environment. These outcomes imply a more efficient use of applied phosphorus, reducing the need for frequent applications and curbing leaching losses that typically accompany soluble phosphate fertilizers. The combination of LDH’s phosphate exchange capacity with biochar’s adsorptive and physico-chemical properties creates a buffering system that can sustain plant-available phosphorus over extended periods, thereby supporting stable growth trajectories and potentially higher yield quality in resource-constrained soils.

Beyond agronomic performance, the study contributes meaningful insights into environmental stewardship and resource sustainability. Phosphorus is a finite, strategically critical nutrient, and global concerns about supply, price volatility, and environmental consequences of fertilizer overuse underscore the need for innovative solutions. Layered double hydroxides provide a reversible phosphate-hosting framework for the recovery and recycling of phosphorus from wastewater, while biochar serves as a durable soil amendment, enhancing soil structure, cation exchange capacity, and microbial habitat. By integrating these components, the P-Mg/Fe-LDH biochar system embodies a circular economy approach that not only optimizes nutrient use efficiency in crop production but also mitigates the risk of nutrient runoff and accumulation in aquatic environments. The study’s findings reinforce the potential of tailored composite materials to bridge the gap between fertilizer effectiveness and environmental protection, contributing to strategies that align with global goals for resource efficiency and food security.

Further exploration is needed to translate these promising results into scalable agricultural practice. Field trials across diverse soil types, climate regimes, and crop species will be essential to validate the robustness and universality of the controlled-release profile observed in greenhouse conditions. Detailed investigations into the long-term fate of LDH-modified biochar in soils, including its impact on soil mineralogy, microbial communities, and potential build-up or degradation over multiple growing cycles, will help elucidate any cumulative environmental effects. Moreover, refining synthesis parameters to optimize loading efficiency, release kinetics, and cost-effectiveness will be critical to commercial viability. Economic analyses should accompany agronomic assessments to compare the total cost of ownership, considering inputs, application frequency, and potential yield benefits versus conventional fertilizers. In summary, this study lays a solid foundation for LDH composite fertilizers that can enhance crop productivity while promoting responsible resource management, providing a viable pathway toward more sustainable agricultural systems that protect both soil health and water quality for future generations.

## CRediT authorship contribution statement

**TS:** Writing-Original draft, Methodology, Investigation, Conceptualization **PMR:** Writing-Original draft, Methodology, Investigation, Conceptualization **RAF:** Supervision, Visualization, Funding Acquisition **JD:** Resources, Supervision, Visualization, Funding Acquisition **JJV:** Resources, Conceptualization **TEM:** Resources, Supervision, Funding acquisition, Project administration

## Acknowledgments

The authors acknowledge the support of the Department of Chemistry, the Department of Plant & Soil Sciences, the Department of Biological Sciences, and the Judy and Bobby Shackouls Honors College at Mississippi State University.

## Data availability

Data used for this study is available at https://github.com/tajindersi/P-LDH-BC

