## Supplement for "Phosphorus-laden Mg/Fe Layered Double Hydroxide Dispersed on Douglas fir Biochar as a Controlled Release Fertilizer and its effect on the growth of bush beans (Phaseolus vulgaris)"

**Table S1.** Average leachate amounts during the BC washing process^[1]^

| Element | Average amount (μg g^-1^) | Element | Average amount (μg g^-1^) |
| --- | --- | --- | --- |
| Fe | 0.8 ± 0.3 | P | 232.6 ± 6.7 |
| Mn | 0.1 ± 0.0 | Li | Not detected |
| V | Not detected | Na | 380.0 ± 12.5 |
| Zr | 0.3 ± 0.0 | Mg | 13.4 ± 0.7 |
| Co | Not detected | K | 5236.5 ± 185.6 |
| Ti | 1.6 ± 0.1 | Ca | 20.6 ± 8.5 |
| Mo | 1.7 ± 0.4 | Ni | Not detected |
| Se | 0.5 ± 0.1 | Zn | Not detected |
| B | 0.1 ± 0.0 | Cd | 0.3 ± 0.0 |
| Si | 4.5 ± 1.3 | Pb | 0.2 ± 0.0 |
| Al | 0.5 ± 0.0 | Ag | Not detected |
| Cr | 0.4 ± 0.0 | Ba | 0.4 ± 0.0 |

**Table S2.** Soil initial nutrient content

| Extractable Nutrient in Pounds Per Acre (ppa) | | Qualitative interpretation as reported in testing results |
| --- | --- | --- |
| Phosphorus (P) | 43 | Low-Medium |
| Potassium (K) | 72 | Low |
| Magnesium (Mg) | 36 | Low-Medium |
| Zinc (Zn) | 0.9 | Low-Medium |
| Organic Sulfur (S) | 99 | Low |
| Calcium (Ca) | 860 | n/a |
| Sodium (Na) | 67 | n/a |

**Table S3.** Randomized block design for pot arrangement including all treatments and replicates

| **9C** | **empty** | **6C** | **3C** | **7C** |
| --- | --- | --- | --- | --- |
| **5C** | **1C** | **2C** | **8C** | **4C** |
| **empty** | **8B** | **9B** | **2B** | **6B** |
| **4B** | **5B** | **1B** | **7B** | **3B** |
| **6A** | **2A** | **8A** | **4A** | **empty** |
| **1A** | **7A** | **3A** | **9A** | **5A** |

**Table S4.** Post-harvest soil nutrient content

| Post-harvest average extractable nutrient levels (Pounds Per Acre) | | | | |
| --- | --- | --- | --- | --- |
| Treatment | P | K | Ca | Mg |
| 1 | 47 ± 2 | 67 ± 9 | 1222 ± 56 | 38 ± 4 |
| 2 | 60 ± 3 | 66 ± 2 | 1183 ± 49 | 36 ± 2 |
| 3 | 78 ± 5 | 52 ± 5 | 1416 ± 97 | 29 ± 2 |
| 4 | 52 ± 4 | 77 ± 7 | 1300 ± 154 | 45 ± 2 |
| 5 | 59 ± 6 | 57 ± 4 | 1293 ± 135 | 31 ± 1 |
| 6 | 79 ± 8 | 58 ± 4 | 1357 ± 66 | 30 ± 3 |
| 7 | 44 ± 9 | 69 ± 9 | 1208 ± 98 | 82 ± 13 |
| 8 | 57 ± 8 | 67 ± 17 | 1210 ± 78 | 78 ± 12 |
| 9 | 82 ± 8 | 50 ± 10 | 1340 ± 82 | 117 ± 19 |

**Table S5.** Average micronutrient uptake from beans

| Average Micronutrient Uptake Per Gram of Beans (μg g^-1^) | | | | | |
| --- | --- | --- | --- | --- | --- |
| Treatment | Zn | Fe | B | Mn | Cu |
| 1 | 42.4 ± 1.4 | 65.8 ± 3.8 | 22.8 ± 2.7 | 19.2 ± 0.4 | 16.5 ± 1.4 |
| 2 | 41.2 ± 0.1 | 69.4 ± 2.3 | 20.9 ± 0.9 | 19.6 ± 1.6 | 14.8 ± 0.5 |
| 3 | 35.0 ± 0.7 | 74.9 ± 6.8 | 22.4 ± 2.3 | 21.6 ± 0.9 | 13.7 ± 0.6 |
| 4 | 35.4 ± 1.3 | 61.4 ± 1.6 | 19.4 ± 0.9 | 18.2 ± 0.8 | 15.3 ± 0.4 |
| 5 | 35.3 ± 1.2 | 70.4 ± 5.7 | 20.4 ± 0.9 | 19.2 ± 1.8 | 15.7 ± 0.3 |
| 6 | 43.9 ± 5.4 | 76.2 ± 1.9 | 23.2 ± 5.2 | 23.7 ± 1.1 | 15.5 ± 0.5 |
| 7 | 39.7 ± 3.9 | 63.5 ± 3.2 | 23.2 ± 0.0 | 18.6 ± 1.3 | 16.0 ± 1.6 |
| 8 | 38.5 ± 3.9 | 70.7 ± 3.2 | 20.3 ± 0.8 | 19.0 ± 0.3 | 14.4 ± 0.9 |
| 9 | 32.3 ± 7.6 | 78.9 ± 2.4 | 19.4 ± 1.3 | 18.3 ± 1.2 | 14.2 ± 0.7 |

**Table S6**. Average micronutrient uptake per gram of remaining plant tissue

| Average Micronutrient Uptake Per Gram of Remaining Plant Tissue (μg g^-1^) | | | | | |
| --- | --- | --- | --- | --- | --- |
| Treatment | Zn | Fe | B | Mn | Cu |
| 1 | 29.7 ± 2.1 | 116.4 ± 1.7 | 30.0 ± 4.7 | 31.4 ± 7.4 | 20.9 ± 1.9 |
| 2 | 24.9 ± 1.7 | 124.6 ± 4.1 | 28.0 ± 3.6 | 28.1 ± 1.7 | 21.5 ± 0.3 |
| 3 | 29.1 ± 1.0 | 152.3 ± 20.5 | 34.3 ± 3.0 | 31.2 ± 2.2 | 19.7 ± 2.0 |
| 4 | 23.5 ± 6.0 | 165.4 ± 10.8 | 28.4 ± 0.2 | 24.5 ± 5.5 | 21.6 ± 0.4 |
| 5 | 23.3 ± 4.1 | 176.1 ± 59.0 | 34.8 ± 0.5 | 26.7 ± 1.0 | 17.4 ± 0.3 |
| 6 | 22.2 ± 6.3 | 151.3 ± 3.1 | 30.0 ± 2.2 | 29.6 ± 1.9 | 22.0 ± 2.4 |
| 7 | 25.4 ± 1.0 | 110.3 ± 2.1 | 28.3 ± 0.0 | 29.3 ± 0.8 | 18.9 ± 0.1 |
| 8 | 25.0 ± 2.2 | 152.8 ± 0.6 | 33.2 ± 0.9 | 28.2 ± 1.0 | 16.7 ± 0.8 |
| 9 | 21.5 ± 0.5 | 144.6 ± 0.0 | 29.6 ± 0.2 | 35.4 ± 9.2 | 17.4 ± 1.5 |

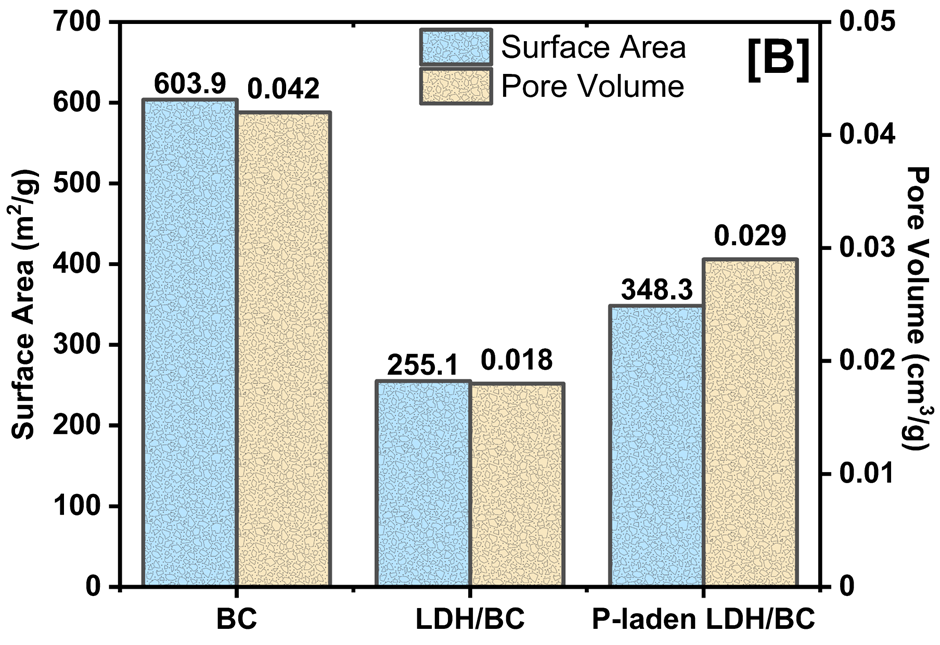

**Fig. S1.** BET surface area and pore volumes of BC, LDH/BC, and P-LDH/BC

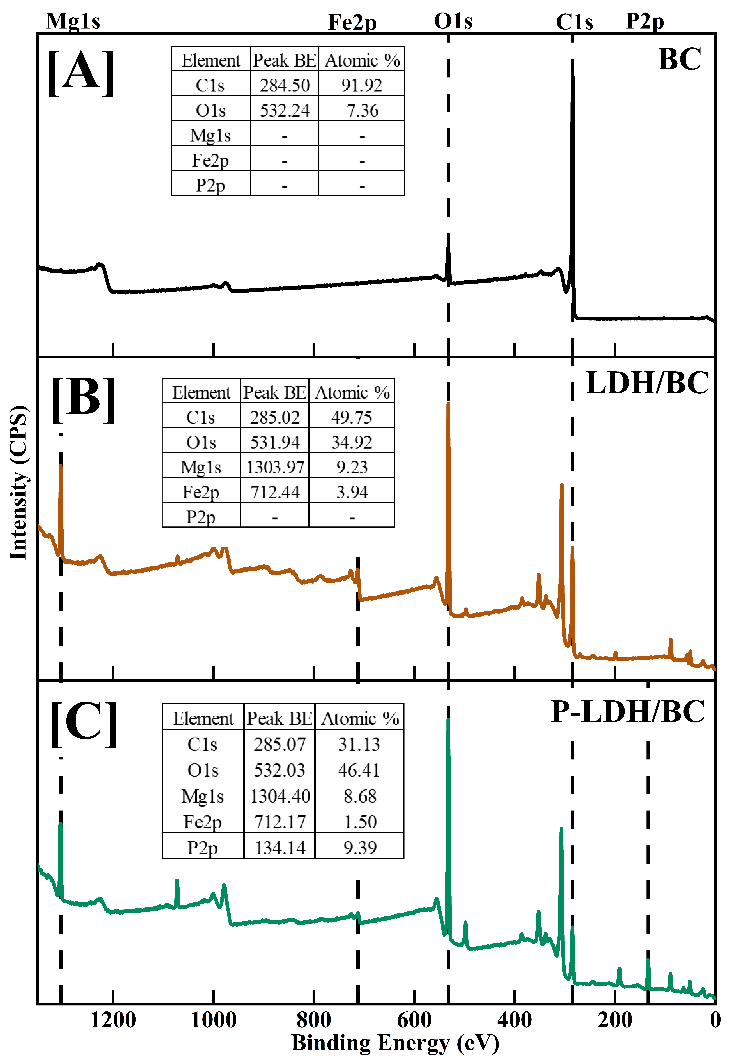

**Fig. S2.** Low-resolution x-ray photoelectron spectroscopy for [A] BC, [B] LDH/BC, and [C] P-LDH/BC

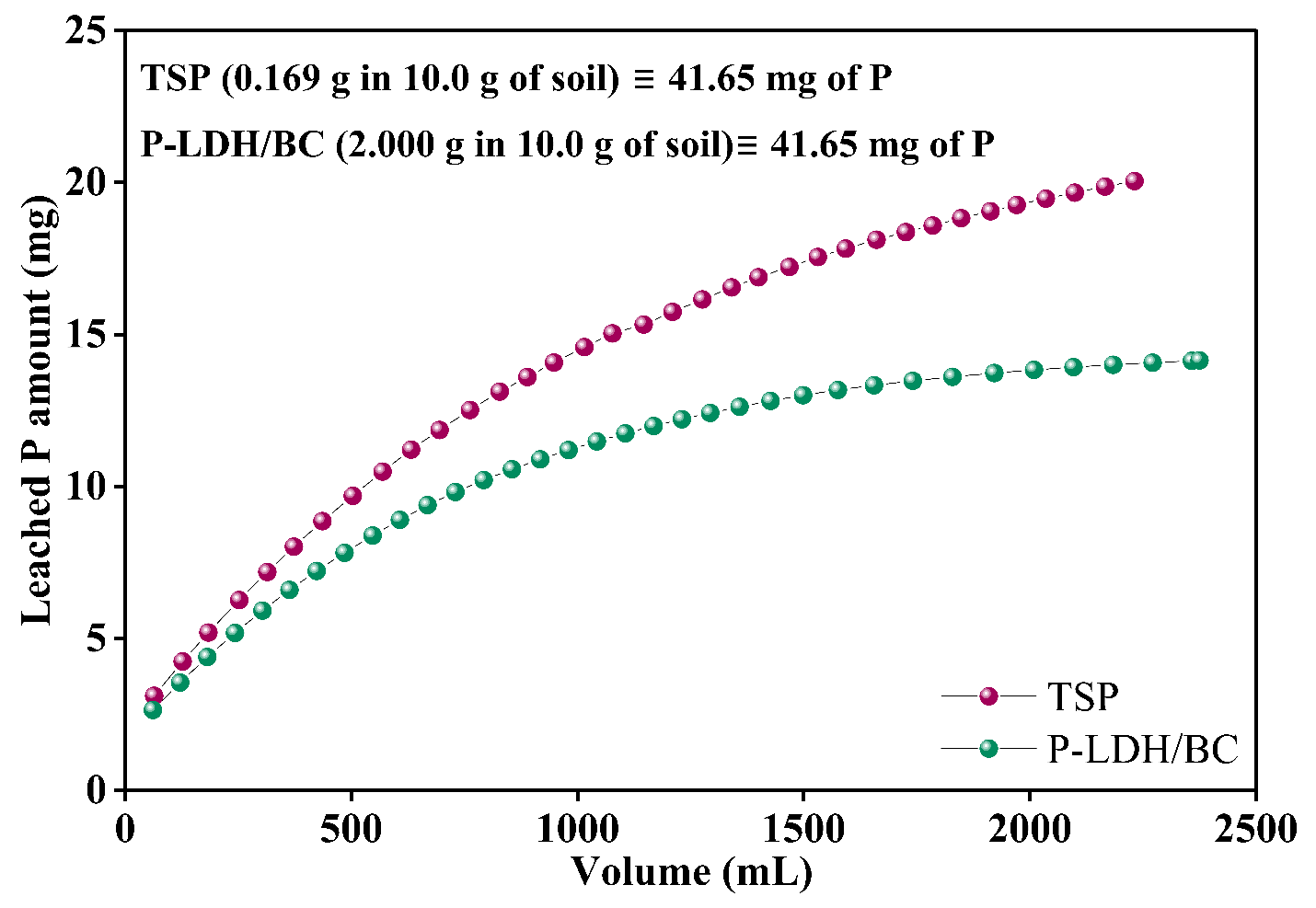

**Fig. S3.** Comparison of P leaching rates of TSP and P-LDH/BC at 1.5 mL min^-1^ flow rate.

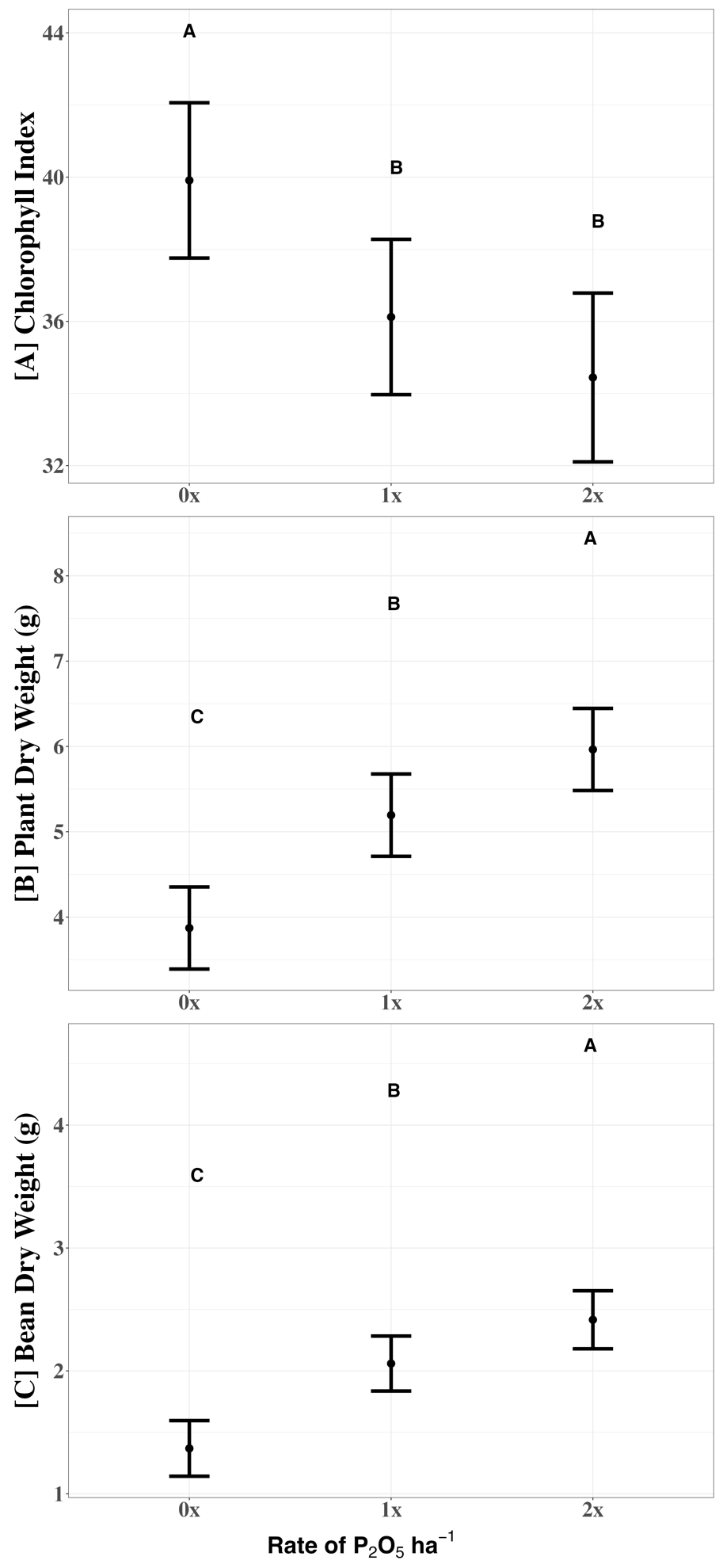
**Fig. S4.** Linear mixed-effect model shows average [A] Chlorophyll Index decreased, [B] Plant dry weight increased, and [C] Bean dry weight increased with increasing rates of P amendment (0x = 0, 1x = 50.44, and 2x = 100.88 kg (P_2_O_5_) ha^-1^) across all treatment types.

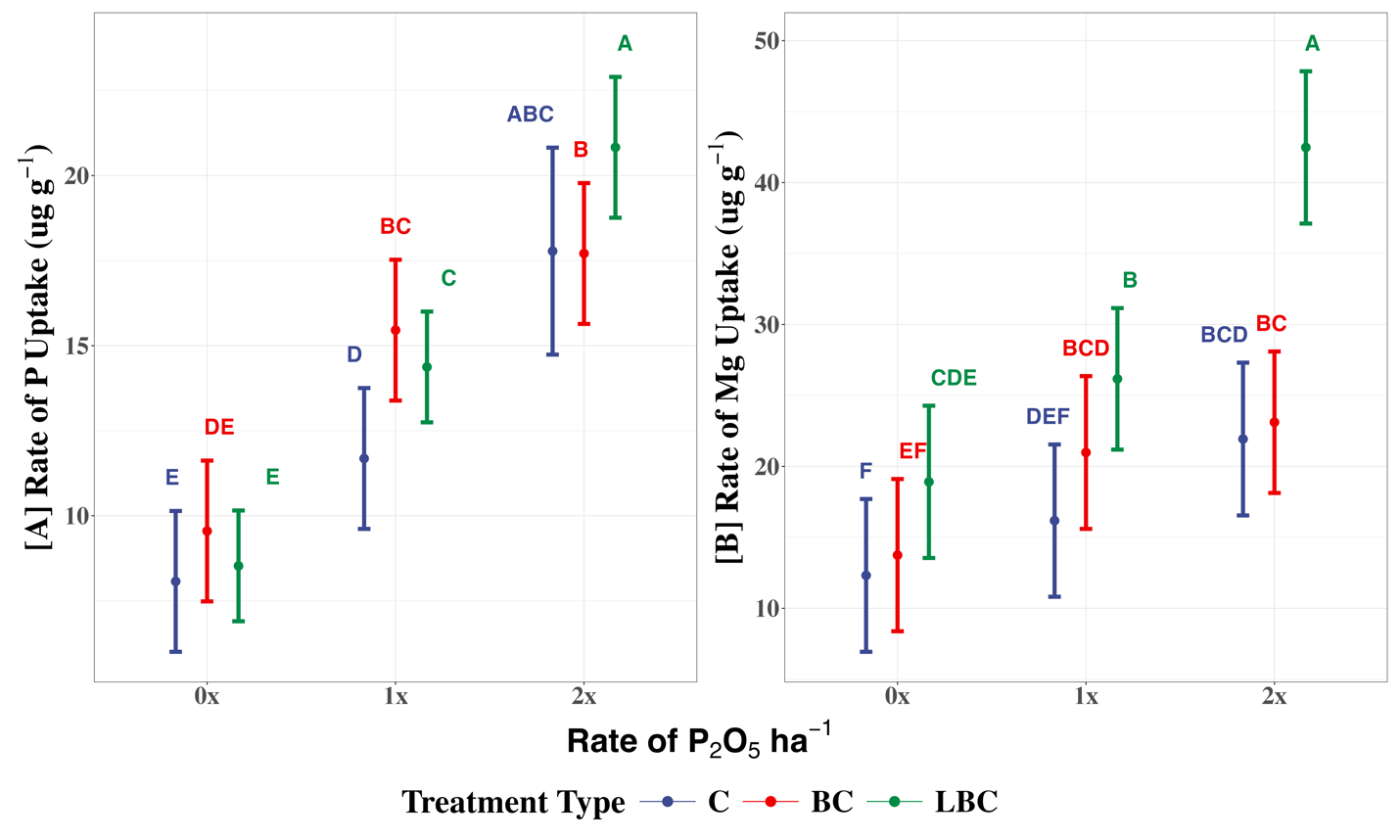

**Fig. S5.** Linear mixed-effect model shows average total [A] P uptake (denoting concentrations) [B] Mg uptake (denoting concentrations) increased with increasing rates of P amendment (0x = 0, 1x = 50.44, or 2x = 100.88 kg (P_2_O_5_) ha^-1^) for all treatment types (C = TSP, BC = BC + TSP, and LBC = P-LDH/BC) and was the highest for plants treated with P-LDH/BC at 100.88 kg (P_2_O_5_) ha^-1^.

1. Values are mean ± SD with 3 replicates
